# Elimusertib outperforms standard of care chemotherapy in preclinical patient-derived pediatric solid tumor models

**DOI:** 10.1101/2022.11.10.515290

**Authors:** Fabian Pusch, Heathcliff Dorado García, Robin Xu, Dennis Gürgen, Yi Bei, Lotte Brueckner, Claudia Röefzaad, Jennifer von Stebut, Victor Bardinet, Rocío Chamorro González, Angelika Eggert, Johannes H. Schulte, Patrick Hundsdörfer, Georg Seifert, Kerstin Haase, Beat Schaefer, Marco Wachtel, Anja A. Kühl, Michael V. Ortiz, Antje M. Wengner, Monika Scheer, Anton G. Henssen

**Author notes:** Corresponding author: Full name: Prof. Dr. med. Anton George Henssen, Email address. These authors contributed equally (Fabian Pusch, Heathcliff Dorado García). These authors jointly supervised this work. Correspondence should be addressed to A.G.H. **Additional information**. Deutsche Forschungsgemeinschaft* (DFG, German Research Foundation) –398299703. Charité 3R, Charité - Universitätsmedizin Berlin. German Cancer Consortium (DKTK). BIH-Charité Clinical Scientist Program funded by the Charité – *Universitätsmedizin Berlin. Research funding by Bayer AG. European Research Council (ERC) under the European Union’s Horizon 2020 research and innovation programme (grant agreement No. 949172). This project received funding by the NIH/CRUK (398299703, the eDynamic Cancer Grand Challenge). Fellowship by “la Caixa” foundation (LCF/BQ/EU18/11650037). Employed by Bayer AG. A.G.H. has received research funding from Bayer AG and is a founder and shareholder of AMZL Therapeutics.

## Abstract

The small molecule inhibitor of ataxia telangiectasia and Rad3-related protein (ATR), elimusertib, is currently being tested clinically in various cancer entities in adults and children. Its preclinical anti-tumor activity in pediatric malignancies, however, is largely unknown. We here assessed the preclinical activity of elimusertib in >40 cell lines and >30 patient-derived xenograft (PDX) models derived from common pediatric solid tumor entities. Detailed *in vitro* and *in vivo* molecular characterization of the treated models enabled the evaluation of response biomarkers. Pronounced objective response rates were observed for elimusertib monotherapy in PDX, when treated with a regimen currently used in clinical trials. Strikingly, elimusertib outperformed standard of care chemotherapies, particularly in alveolar rhabdomysarcoma PDX. Thus, elimusertib has strong preclinical anti-tumor activity in pediatric solid tumor models, which may translate to clinically meaningful responses in patients.

**Statement of translational relevance:** Elimusertib is a small molecule inhibitor of ATR. ATR inhibitors have shown promising results as anticancer agents in adult cancers, but there is limited information on their effectiveness in pediatric solid tumors. Using a cohort of 32 patient-derived xenografts from pediatric solid tumors, we here evaluated the therapeutic potential of elimusertib *in vivo*. Elimusertib reduced tumor volume growth in all samples. Elimusertib had very limited toxicity and was potent even in tumors with preexisting chemoresistance. Our preclinical data indicates that elimusertib is a safe and potent therapeutic option for pediatric solid tumors. This data may serve as a rationale for the development of pediatric clinical trials for ATR inhibitors.

## Introduction

Pediatric cancers are rare but represent a leading cause of death in children (1). Currently, pediatric solid tumors are treated with a histology-specific and risk-stratified combination of surgery, radiotherapy, and chemotherapy. Despite steady improvements in the survival rate of childhood cancers over the last several decades (2), cures remain unacceptably low for many high risk pediatric solid tumors. Even for those who are ultimately cured, the aggressive multi-modality approaches are frequently associated with severe long-term morbidities (3). As a result, there is an urgent need to identify novel therapeutic approaches, which leverage specific tumor vulnerabilities.

Compared to adult cancers, which often demonstrate high numbers of mutations accumulated over a lifetime, pediatric tumors generally arise during developmental windows in a tissue-context specific manner, often harboring only few mutational drivers and a low mutational burden (4). A common feature among pediatric solid tumors is the presence of fusion oncoproteins, which emerge as a result of chromosomal aberrations (5). Additionally, intra- and extrachromosomal oncogene amplifications are frequent in certain pediatric solid tumors, such as in neuroblastoma, where *MYCN* amplifications, often occurring on ecDNA, are a predictor for poor prognosis (6-10). Both gene amplifications and fusion oncoproteins are hard to therapeutically target directly, particularly when affecting transcription factors, which has hampered the development of selective therapies in these tumor entities.

Genomic instability is a hallmark of cancer cells (11), which has recently been shown to be therapeutically actionable (12). The extreme proliferation rate in cancer cells, in part induced by fusion oncoproteins and oncogene amplifications, can result in delays or errors in the DNA termed replication stress (13-15). In response to the damaged DNA, cells have intricate mechanisms to recognize and repair lesions while ensuring that the cell cycle is halted, termed the DNA damage response (DDR). The DDR is mainly regulated by three kinases: ataxia telangiectasia mutated (ATM), ataxia telangiectasia- and Rad3-related (ATR), and DNA-dependent protein kinase catalytic subunit (DNA-PKcs) (16). Even though they have similar protein sequences, and their targets overlap, it is widely accepted that they respond to different stimuli (17). While ATM and DNA-PKcs are mostly activated after double strand breaks (DSBs), ATR responds primarily to replication stress-associated DNA damage, which often involves single stranded DNA intermediates (18,19). Because ATR is activated in response to replication stress, it has been suggested that cancers depend on ATR more strongly than non-transformed cells to tolerate high levels of replication stress (20,21). These findings have fueled the interest to test ATR inhibitors as a therapeutic option in cancer, particularly in tumors with high replication stress. Some biomarkers for predicting ATR inhibitor response have been put forward, e.g. ATM loss, TP53 loss, MYC overexpression, CDC25A overexpression, PGBD5 expression and fusion oncoproteins such as EWS-FLI1 and PAX3-FOXO1, which increase sensitivity to ATR inhibitors (22-30) and are currently considered in clinical trial design (NCT04095273, NCT03188965, NCT03682289, NCT04170153, NCT04576091, NCT04535401, NCT04657068, NCT05338346, NCT04616534, NCT04514497, NCT05071209). How most pediatric solid tumor entities may benefit from ATR inhibitor treatment is difficult to predict, as detailed preclinical information is currently missing.

Here we profiled the anti-tumor effects of the ATR inhibitor elimusertib (also known as BAY 1895344) *in vitro* and in a cohort of PDXs from pediatric solid tumors. In order to create a solid basis for future clinical trial designs, we compared the effects of elimusertib to those of first-line standard-of-care (SoC) chemotherapeutics. We demonstrate that monotherapy with elimusertib has most pronounced antiproliferative effects in models of alveolar rhabdomyosarcoma and neuroblastoma, and identify specific molecular alterations that may predict response to elimusertib. These findings highlight a potential therapeutic role for ATR inhibition in a subset of childhood solid tumors and provide a basis to accelerate the translation into meaningful clinical applications.

## Materials and Methods

### Study design

The purpose of this study was to examine the effects of ATR inhibition in preclinical models of pediatric solid tumors and identify potential biomarkers to select patients that could benefit from a treatment with the ATR inhibitor elimusertib. We first determined the inhibitory activity of the elimusertib in cell models, and compared these cells based on known determinants of ATR inhibition sensitivity, as well as the presence of oncogenes which increase the level of replication stress. We analyzed the effects of elimusertib treatment on cell cycle control and genomic instability. All *in vitro* experiments were performed following the guidelines proposed by Carola A.S. Arndt for pediatric tumors (31). In the study, five to eight cell lines were used per disease, for which we validated the expression of the target genes and included the elimusertib IC_50_ after 72h. Outliers were not excluded unless technical errors were present. For *in vivo* testing, sample size was decided based on previous experience with the models. Animals euthanized before the end of the experiment, due to excessive tumor growth or loss of body weight, were included in the analysis. The researchers and patients were not blinded during the experiments.

### Reagents

All reagents were obtained from Carl Roth (Karlsruhe, Germany) unless otherwise indicated. Elimusertib was provided by Bayer AG (Leverkusen, Germany). Elimusertib was dissolved in dimethyl sulfoxide (DMSO) and stored at 10 mM concentrations at -20 °C until further use.

### Cell culture

All neuroblastoma and Ewing sarcoma cell lines were kindly provided by Prof. J.H. Schulte (Charité). Rh41, Kym1 and Rh18 cells were a kind gift from Prof. Simone Fulda (Kiel, Germany). The remaining human tumor cell lines were obtained from the American Type Culture Collection (ATCC, Manassas, Virginia). All rhabdomyosarcoma and all Ewing’s sarcoma cell lines, as well as RPE and BJ cell lines were cultured in Dulbecco’s Modified Eagle’s Medium (DMEM, Gibco, Thermo Fisher Scientific, Waltham, Massachusetts, USA) supplemented with 10% fetal calf serum (Thermo Fisher) and penicillin/streptomycin (Gibco, Thermo Fisher Scientific). All neuroblastoma cell lines were cultured in Roswell Park Memorial Institute (RPMI)-1640 (Gibco, Thermo Fisher Scientific) supplemented with 10% fetal calf serum and penicillin/streptomycin. Twice per week, cells were washed with phosphate-buffered saline (PBS), incubated in 0.05% Trypsin-EDTA (1x) (Gibco, Thermo Fisher Scientific) for five minutes, resuspended in culture medium, sedimented at 500 g for 5 minutes and a fraction was cultured in fresh media. Cells were kept in culture for a maximum of 30 passages. Resuspended cells were counted by mixing 1:1 with 0.02 % trypan blue in a BioRad (Hercules, CA, USA) TC20 cell counter. The absence of *Mycoplasma sp*. contamination was determined using a Lonza (Basel, Switzerland) MycoAlert system.

### Cell viability

Cell viability was assessed using CellTiter-Glo (Promega, Madison, Wisconsin, USA). Briefly, for CellTiter-Glo measurement, 1,000 cells were seeded in white, flat-bottom, 96-well plates (Corning, Corning, NY, USA). After 24 hours, drugs were added to the medium and cells were incubated for 72 hours. CellTiter-Glo luminescent reagent was added according to the manufacturers protocol, and the luminescence signal measured on a Glowmax-Multi Detection System (Promega).

### Western Immunoblotting

Whole-cell protein lysates were prepared by lysing cells in Radioimmunoprecipitation assay buffer (RIPA) supplemented with cOmplete Protease inhibitor (Roche, Basel, Switzerland) and PhosphStop (Roche). Protein concentrations were determined by bicinchoninic acid assay (BCA, Thermo Fisher). 10 μg of protein were denatured in Laemmli buffer at 95 °C for 5 minutes. Lysates were loaded onto 16%, or 10% Tris-Glycin (Thermo Fisher) gels for gel electrophoresis depending on the protein sizes of interest. Proteins were transferred onto Polyvinylidenfluorid (PVDF) membranes (Roche), blocked with 5% dry milk or 5% bovine serum albumin for 1 hour and incubated with primary antibodies overnight at 4°C, then secondary antibodies for 1 hour at room temperature. Chemiluminescent signal was detected using Enhanced chemiluminescence (ECL) Western Blotting Substrate (Thermo Fisher) and a Fusion FX7 imaging system (Vilber Lourmat, Marne-la-Vallée, France). Quantification was performed with ImageJ.

### Immunofluorescence staining

Cells were grown at the desired confluency on glass slides with an 8 well flexiPERM silicone grid (Sarstedt, 94.6032.039) for 24h and directly processed (for R-loop quantification) or treated with 20 nM elimusertib for 48 h (micronuclei quantification). Cells were washed with PBS three times and fixed for 10 minutes with 3.7 % paraformaldehyde, washed with PBS three times and permeabilized with PBS containing 0.1% Triton-X100. For R-loop immunofluorescence cells were blocked for 30 minutes with 10% FCS in PBS-T (0.2% Tween-20 in PBS), incubated overnight at 4^a^C with the primary antibody (Anti-DNA-RNA Hybrid Antibody, clone S9.6; Merck Millopore MABE1095), washed three times with PBS-T (0.05% Tween-20 in PBS), incubated for 1 hour in the dark at room temperature with the secondary antibody (Dianova, 715-096-150). After removal of the 8 well silicone grid, the glass slide was washed three times with PBS-T (both R-loop and micronuclei quantification). The glass slide was covered with DAPI-containing mounting media (Vectashield, Vec-H-1000) and mounted with a cover slip. Cells were imaged using an ECHO Revolve microscope and quantified using ImageJ.

### Fluorescence-activated cell sorting (FACS)

Cells were grown in the presence of drug or vehicle (DMSO) for 72h prior to sample preparation for flow cytometry. For cell cycle analysis, cells were incubated with 5-Ethynyl-2’
s-deoxyuridine (EdU) for 2 hours right before fixation and fluorescent labeling, following the instructions provided in the kit Click-IT EdU Alexa Fluor 488 Flow Cytometry Assay kit (Thermo Fisher). For DNA damage analysis, terminal deoxynucleotidyl transferase dUTP nick end labeling (TUNEL) was performed using the APO-BrdU TUNEL Assay Kit (Thermo Fisher), according to the manufacturer’s descriptions. Stained cells were measured on a BD LSR Fortessa flow cytometer (BD Biosciences, Franklin Lakes, NJ, USA) and analyzed using FlowJo (v 10.8.1).

### Patient-derived xenograft (PDX) treatment

The establishment of PDX models was conducted as previously described (32) in collaboration with Experimental Pharmacology & Oncology GmbH (EPO, Berlin, Germany). All experiments were conducted according to the institutional animal protocols and the national laws and regulations. Tumor fragments from rhabdomyosarcoma patients were transplanted into either Crl:NMRI-*Foxn1*^*nu*^ mice (Charles River, Wilmington, MA, USA) or NOD.Cg-Prkdc^scid^ Il2rg^tm1Sug^/JicTac mice (Taconic, Rensselaer, NY, USA). Tumor growth was monitored with caliper measurements. Tumor volume was calculated with the formula length x width^2^ / 2. PDX were serially transplanted in mice at least three times prior to the experiments. Mice were randomized into four groups with at least 3 mice to receive treatment. For the elimusertib study, mice were administered 40 mg/kg body weight on a 3 days on/4 days off regime twice daily (orally). Elimusertib was dissolved in 60% polyethylene glycol 400, 10% ethanol and 30% water to a 4mg/ml solution, the same solution without compound was used as vehicle control. Mice were sacrificed by cervical dislocation once the tumor volume exceeded 1.500 mm^3^ or body weight loss was higher than 20%.

### Immunohistochemistry stainings

Paraffin sections of 1 μm thickness were cut, dewaxed and subjected to a heat-induced epitope retrieval step. Endogenous peroxidase was blocked by hydrogen peroxide prior to incubation with anti-Ki67 (clone D2H10, Cell Signaling Technologies), anti-Histone H3-S10 (polyclonal rabbit, Abcam #47297) or anti-γH2AX (polyclonal rabbit, Abcam #229914) followed by incubation with EnVision+ HRP-labelled polymer (Agilent). For visualization, 3,3’-diaminobenzidine (DAB) as chromogen was used. For detection of cleaved caspase3, anti-clCasp3 (clone 5A1E, Cell Signaling Technologies) was used followed by incubation with secondary antibody (biotinylated donkey anti-rabbit) and alkaline phosphatase-labelled streptavidin (Agilent). RED was used as chromogen (Agilent). Nuclei were stained with hematoxylin (Merck) and slides were coverslipped in glycerol gelatine (Merck). Multispectral images were acquired using a Vectra^®^ 3 imaging system (Akoya Biosciences). The QuPath software (version 0.3.2) was used for cell segmentation as well as quantification.

### Cell line and PDX genomic analysis

Cell line mutation data was obtained from the online public dataset DepMap (https://depmap.org/portal/, packages Copy Number Public 21Q2 and Mutation Public 21Q2). WGS, WES and RNA sequencing from the PDX samples was performed using NEBNext Ultra II FS DNA library Kit for Illumina (New England Biolabs), SureSelectXT HS Target Enrichment System for Illumina Paired-End Multiplexed Sequencing Library For Illumina Multiplexed Sequencing Platforms (Agilent), and TruSeq Stranded mRNA Library Prep (New England Biolabs), respectively, following the protocol provided by the manufacturers. Oncoplots were drawn using the R package maftools (v 2.12.0).

### Statistical analysis

All statistical tests were done using GraphPrism9 or R.

### Data availability

The data generated in this study are available upon request from the corresponding author. Restrictions apply to the availability of data that does not comply with patient privacy requirements.

## Results

### Elimusertib treatment affects survival of pediatric solid tumor cell lines

To study the therapeutic potential of elimusertib inhibition in pediatric solid tumors, we treated 41 cell lines derived from several pediatric tumors, including Ewing’s sarcoma (EWS), alveolar (ARMS) and embryonal rhabdomyosarcoma (ERMS) and high-risk neuroblastoma with and without *MYCN* amplification (MNA NB vs. NMNA NB), with the ATR inhibitor elimusertib and measured their survival over time (Fig. 1a-e). Cells showed a wide range of response, with inhibitory 50% concentrations (IC_50_) values ranging from 2.687 to 395.7 nM (Extended Data Table 1). These concentrations are well below plasma concentrations achievable in human patients (33), suggesting that elimusertib may exert similar anti-tumor effects *in vivo*. Compared to non-transformed cell lines BJ and RPE cells, elimusertib inhibited cell viability at lower concentrations in most cancer cell lines (Fig. 1f). In line with previous reports testing other ATR inhibitors (24,26,29), cell lines derived from Ewing sarcoma, *MYCN*-amplified neuroblastoma and alveolar rhabdomyosarcoma were (significantly) more sensitive to ATR inhibition than control cell lines, suggesting a therapeutic window may exist for elimusertib in these pediatric solid tumors.

**Figure 1.**
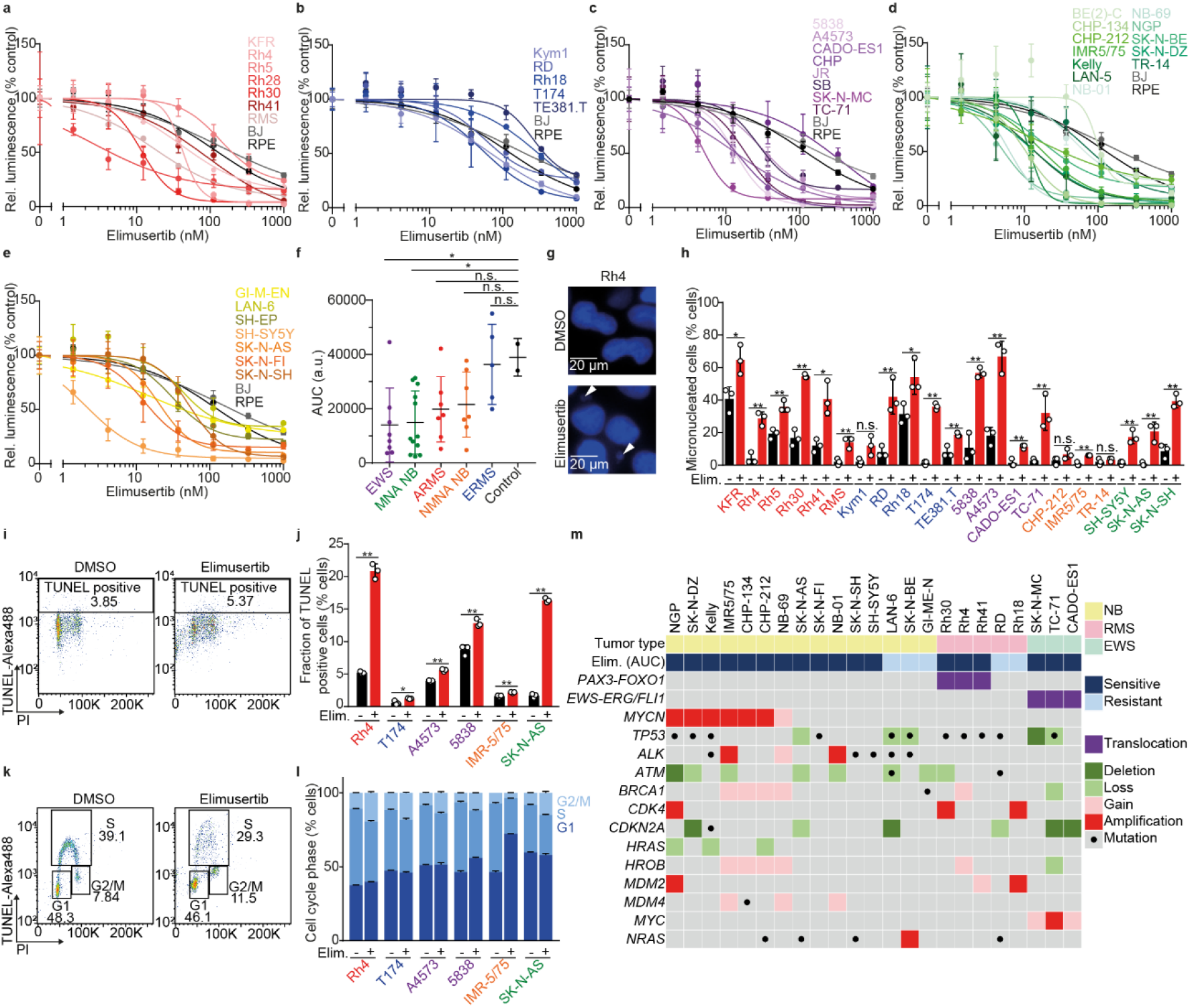
Elimusertib shows anti-tumor activity in a broad spectrum of pediatric cancer cell lines. **(a-e)** Dose-response curves of the cell viability for ARMS (a), ERMS (b), EWS (c), MNA NB (d) and NMNA NB cell lines (e) treated with the ATR inhibitor elimusertib compared to non-cancer cell lines BJ and RPE (*n* = 3; 50% inhibitory concentrations, IC_50,_ and area under the curve, AUC, values are listed in Extended Data Table 1). **(f)** AUC corresponding to the graphs in (a-e) (unpaired, two-sided Student’s t test, *P* = 0.0410, 0.0165, 0.0761, 0.0992, 0.8260 for EWS vs Control, MNA NB vs Control, ARMS vs Control, NMNA NB vs Control, ERMS vs Control, respectively). **(g)** Representative photomicrographs of micronuclei (white arrow) in cells treated with elimusertib. **(h)** Fraction of micronucleated cells after treatment with elimusertib (20 nM) for 72h (*P* = 0.0242, 0.0014, 0.0033, 0.0002, 0.0108, 0.0065, 0.520, 0.0061, 0.0312, 1.30×10^−5^, 0.0072, 0.0008, 0.0014, 0.0026, 0.0088, 0.1448, 0.0013, 0.3740, 0.0030, 0.0042, 0.0008, respectively; n = 3, with 50 cells per replicate). **(i)** Representative gating for TUNEL labeling in 5838 cells. **(j)** Quantification of TUNEL signal in a set of pediatric solid tumor cell lines treated with elimusertib (20 nM) for 72h. (*P* = 2.08×10^−5^, 0.0232, 0.0002, 0.0018, 0.0045, 6.38×10^−7^, respectively; n = 3). **(k)** Representative gating for EdU and PI co-staining in 5838 cells. **(l)** Quantification of the fraction of cells in each cell cycle phase in a set of pediatric solid tumor cell lines after elimusertib treatment (20 nM) for 72h (n = 3; unpaired, two-sided Student’s t test; error bars represent standard deviation). **(m)** Table of mutations (incl. translocations, single nucleotide variants, copy number alterations) affecting genes associated with ATR inhibitor sensitivity in a subset of cell lines tested.

### Elimusertib treatment leads to DNA damage in pediatric solid tumor cell lines

ATR is a key regulator of replication stress-induced DNA damage (18,34,35). To investigate the effects of ATR inhibition in pediatric cancer cell lines, we measured DNA damage accumulation in response to elimusertib treatment in a subset of cell lines. Micronucleation is an indicator of genomic instability (36). In response to elimusertib, cell lines showed higher rates of micronucleation (Fig. 1g-h), indicating the presence of DNA damage. Co-staining with TdT-dependent UTP nicked-end labelling (TUNEL) and propidium iodide indicated an increase in the fraction of cells with fragmented DNA in cells incubated with elimusertib, suggesting an accumulation of unrepaired damaged DNA and apoptotic DNA fragmentation (Fig. 1i-j), which is in line with previous reports (26,29,33,37,38). Because ATR is crucial for the intra-S and G2/M checkpoint activation (39-41), we examined cell cycle progression in response to elimusertib. We pulse-labelled replicating DNA with 5-Ethinyl-2’-Desoxyuridin (EdU) and stained all DNA with propidium iodide in cells incubated in the presence of elimusertib. In all cell lines tested, elimusertib led to a reduction in the fraction of cells in S-phase, consistent with a repression of the intra-S checkpoint. In all cell lines but one (IMR-5/75), we observed an increase in cells in G2/M (Figure 1k-l). To assess whether cells accumulated in mitosis, consistent with a G2/M checkpoint suppression, we measured Histone 3 phosphorylation at Serine 10, a marker specific for mitosis (42). After incubation in the presence of elimusertib, we did not observe a consistent increase in IMR-5/75 (neuroblastoma) and A4573 (Ewing sarcoma) cells, suggesting cell context dependent cell cycle disruption in response to elimusertib (Extended Data Fig. 1a-b). We next evaluated the effect of elimusertib on replication stress by measuring RPA32 T21 phosphorylation, in cells incubated with elimusertib. RPA32 phosphorylation, a marker of single-stranded DNA, was increased in response to elimusertib (Extended Data Fig. 1a-b). Taken together, this suggests that elimusertib prevents repair of replication stress-associated DNA damage, resulting in further genomic instability and then ultimately apoptosis in these pediatric solid tumor cell line models.

### Fusion oncoprotein expression and high MYCN levels are associated with elimusertib sensitivity

Because ATR is key in repairing replication stress-induced DNA damage, we tested whether cell lines with varying levels of ATR-mediated replication stress response signaling would differ in their sensitivity to elimusertib. For this purpose, we assessed the abundance of R-loops, a nucleic acid structure consisting of and RNA:DNA hybrid and single stranded DNA which has been implicated in genomic instability as well as replication stress and is being discussed as mediator for treatment susceptibility in cancer (43,44). In contrast to previous reports, no positive correlation was observed between the abundance of R-loops and elimusertib sensitivity (Extended data fig. 2a-c). Sensitivity to ATR inhibitors can be influenced by genetic aberrations frequent in cancers, such as TP53 or ATM loss, PGBD5, MYC(N) expression, or fusion oncoproteins such as EWS-FLI1 and PAX3-FOXO1 (22,24-27,29,45). We assessed the presence of frequent genetic alterations in pediatric tumors (46) as well as markers that cause genetic vulnerability to ATR inhibition (22,25,27,28,47,48) in our cell lines using publicly available datasets (49). In line with previous reports (28), the presence of *MYCN* amplifications, both on ecDNA or as homogenously staining regions (50,51), in NB cell lines, expression of fusion oncoproteins such as EWS-FLI1 or PAX3-FOXO1 (25,29) and TP53 deficiency (22) were associated with higher elimusertib sensitivity (Fig. 1m). Thus, the presence of known biomarkers of ATR inhibitor sensitivity is also associated with elimusertib sensitivity in pediatric tumor cell lines and may be suitable for patient selection in current and upcoming clinical trials.

### A preclinical trial of elimusertib in patient-derived xenografts demonstrates clinically relevant response in a large subset of pediatric solid tumors

Encouraged by the results obtained *in vitro*, we sought to test the preclinical anti-tumor activity of elimusertib *in vivo* in mice harboring patient-derived xenograft models (PDX) of pediatric solid tumors (Fig. 2a). We selected a cohort of PDX derived from 8 EWS, 4 ERMS, 7 ARMS, 4 MNA-NB, 5 NMNA-NB, 3 osteosarcomas (OS) and one CIC-DUX fusion gene expressing undifferentiated sarcoma. Within each entity, the cohort comprised various sites of origin, primary or relapse status, histopathological gradings and clinical stagings (Extended Data Table 2). In total, we treated 195 mice (median 3 mice per PDX model and treatment arm) and 32 PDX models derived from patients treated at the Charité – Universitätsmedizin Berlin (Xu et al., currently under consideration elsewhere) and the University Children’s Hospital, Zurich (52). Some PDX were derived from the same tumors but collected before and after treatment (EWS_3a and EWS_3b) or sequential relapses (ERMS_2a, ERMS_2b and ERMS_2c) (Extended Data Table 2). In order to closely mirror the setup of a clinical trial, we treated mice using the same regimen currently used in clinical trials, i.e. elimusertib at 40 mg/kg body weight twice daily per oral gavage, on a 3-days on/4-days off schedule for 28 days (Fig. 2a). According to the Response Evaluation Criteria in Solid Tumours (RECIST) (53,54), two of the PDX models achieved a complete response (CR), two PDX had a partial response (PR), 14 PDX were considered as stable disease (SD), and 16 PDX were classified as progressive disease (PD, Fig. 2b-d). In all cases, single agent elimusertib treatment was sufficient to significantly delay tumor growth, compared to vehicle-treated control mice (Extended Data Fig. 3a-af). Consistent with our previous work using AZD6738 (29) mice harboring PDX derived from ARMS showed the most pronounced response, with only one out of the seven ARMS PDX models classified as progressive disease after elimusertib treatment (Extended Data Fig. 3a-g) ERMS (Extended Data Fig. 3h-k) and MNA NB PDX (Extended Data Fig. 3w-aa) also showed a good response, with only one and two models with progressive disease, respectively. Interestingly, the ERMS model derived from a later relapse showed a better response than the models derived from the same patient at an earlier timepoint (ERMS_2a and EMRS_2b, respectively; Fig. 2b-c, Extended Data Fig. 3i-k), implicating that treatment-associated tumor evolution may have enhanced ATR inhibitor sensitivity. Toxicity, assessed by body weight loss over time, was minimal during treatment, indicating a good tolerability of the drug in the given regimen (Extended Data Fig. 4a-af). Together, elimusertib monotherapy has clinically relevant anti-tumor activity in pediatric solid tumor models.

**Figure 2.**
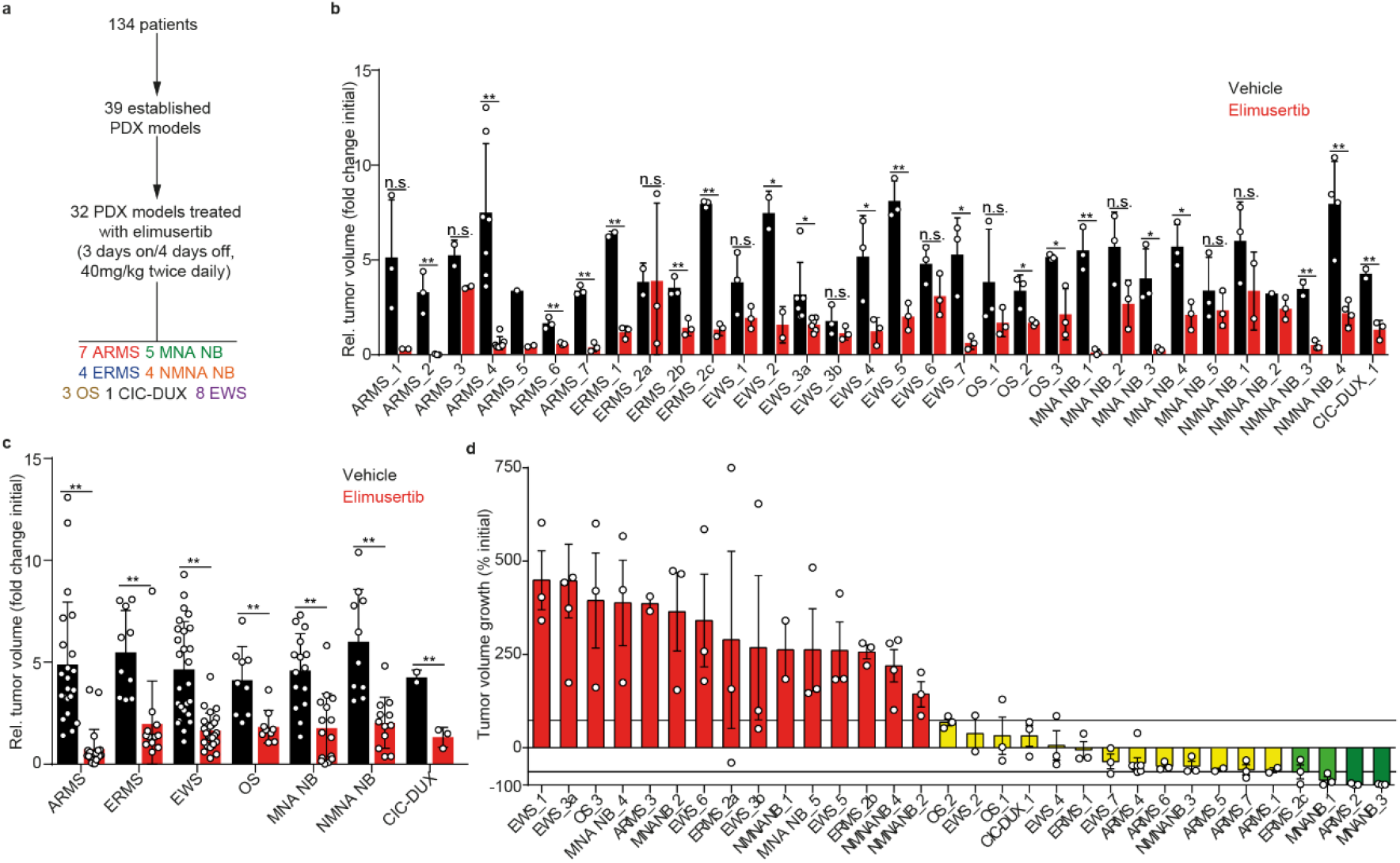
Elimusertib treatment induces heterogeneous response in a large cohort of patient-derived xenografts of pediatric solid tumors. **(a)** Schematic representation of the preclinical study in PDX models. A total of 39 PDX models were established from 134 patients. 32 of those PDXs received 40 mg/kg body weight elimusertib twice daily per oral gavage, on a 3 days-on/4 days-off schedule. **(b)** Dot plot showing the relative tumor volume at the end of the treatment for all PDXs treated with elimusertib or vehicle control (*n* and *P* values are listed in Extended Data Table 3). **(c)** Dot plot showing the relative tumor volume at the end of the treatment for all tumor entities treated with elimusertib or vehicle control (*n* and *P* values are listed in Extended Data Table 3). **(d)** Waterfall plot representing tumor volume change in mice receiving elimusertib. Tumors were classified according to the RECIST criteria(54) as progressive disease (red), stable disease (yellow), partial response (light green) and complete response (dark green). For statistical comparison an unpaired, two-sided Student’s t test was performed; error bars represent standard deviation.

### Elimusertib treatment extends progression-free survival in pediatric solid tumor models

In order to further evaluate the preclinical activity of elimusertib, we assessed the progression-free survival (PFS) of PDX after elimusertib treatment. Overall, elimusertib extended the median PFS from 7 to 20 days across PDX models from different tumor entities (Fig. 3a). The most pronounced extension of PFS was observed for ARMS (Fig. 3b, median PFS from 9 days to the end of experiment), followed by ERMS (Fig. 3c, median PFS from 5 to 26 days). Median PFS increased from 7 to 14 days for EWS (Fig. 3d), from 6 to 12 days for MNA NB (Fig. 3e), 7 days to 17 for NMNA NB (Fig. 3f), 9 to 20 days for OS (Fig. 3g) and 5 to 12 days for the CIC-DUX model (Fig. 3h). Furthermore, elimusertib prolonged overall survival across PDX from all tumor entities with a median overall survival of 19 days vs. 31 days in the untreated and elimusertib-treated group, respectively (Extended Data Figure 5a). For some tumor entities, such as ARMS, ERMS, NMNA NB, and OS, the overall survival rate in the treatment group was significantly higher than the control group at 30 days, exceeding 75% overall survival (Extended data Fig. 5b, c, f, g). MNA NB and EWS also showed significantly prolonged overall survival, whereas the overall survival of the CIC-DUX models was not statistically significant (Extended data figure 5d, e, h). Thus, elimusertib monotherapy delays tumor growth, which results in pronounced increases in PFS and overall survival in diverse pediatric solid tumor models.

**Figure 3.**
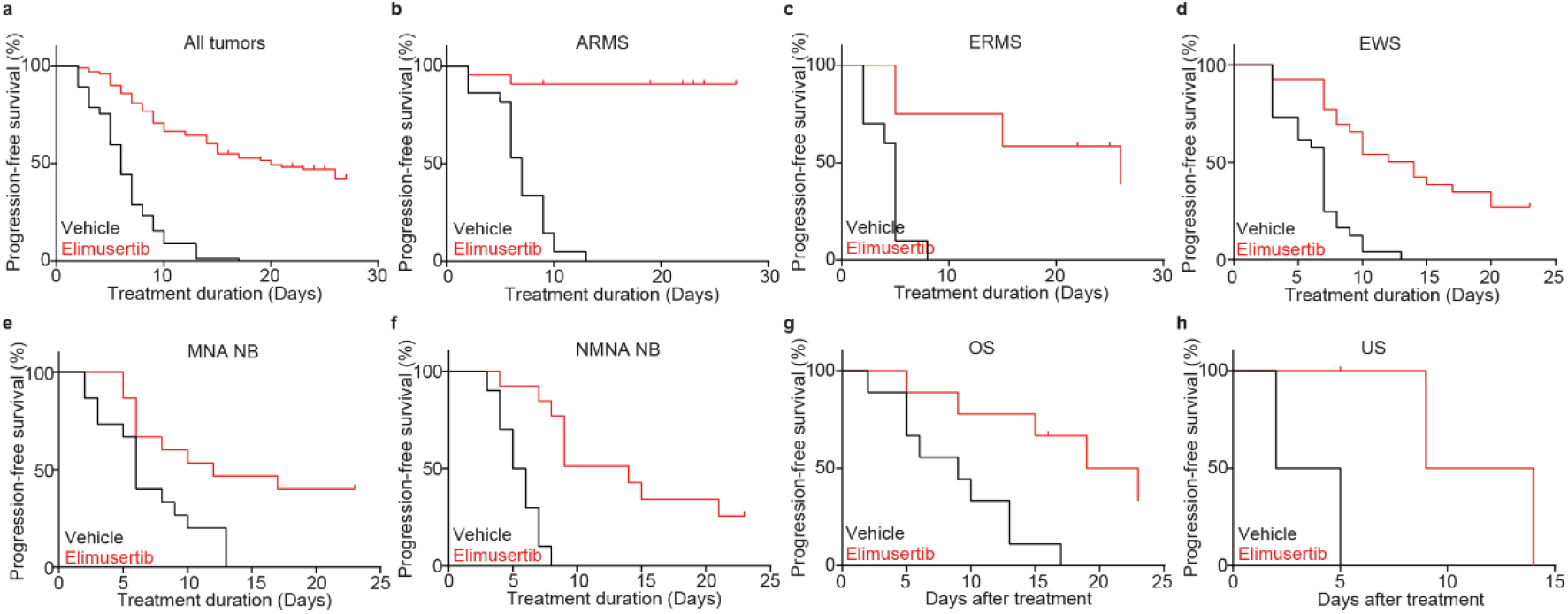
Elimusertib treatment extends the progression-free survival of preclinical pediatric solid tumor models. **(a-h)** Kaplan Meier curves showing the progression-free survival, defined as time until the tumor was classified as progressive disease, PD, according to the RECIST criteria, in mice treated with elimusertib (red) or vehicle (black), across tumor types (a, *n*_*total*_ = 195, *P* < 0.0001), ARMS (b, *n*_*total*_ = 44, *P* < 0,0001), ERMS (c, *n*_*total*_ = 22, *P* = 0.0001), EWS (d, *n*_*total*_ = 53, *P* = < 0.0001), MNA NB (e, *n*_*total*_ = 30, *P* = 0.0064), NMNA NB (f, *n*_*total*_ = 23, *P* < 0.0001), OS (g, *n*_*total*_ = 18, *P* = 0.0033) and CIC-DUX sarcoma (h, *n*_*total*_ = 5, *P* = 0.0389). Log-rank tests were performed for statistical comparison.

### Reduced proliferation rate in pediatric solid tumor PDX after elimusertib treatment represents a putative response biomarker

To characterize the effect of elimusertib treatment on PDX, we performed immunohistochemical (IHC) staining of molecular markers of cell proliferation, DNA damage and apoptosis in 21 of the 32 PDX models at the end of elimusertib treatment (Extended Data Fig. 6, 7, 8, 9 & 10; Extended Data Table 4, 5 & 6). Baseline expression of these markers was not associated with differences in elimusertib response (Extended Data Fig. 11a, c-d). Only high pre-treatment Histone H3 phosphorylation (pHH3) expression, indicative of mitotic cells, was slightly associated (not statistically significant) with good PDX response (Extended data fig 11b). The fraction of Ki-67 positive cells, an indicator of proliferating cells, in PDX was significantly lower in elimusertib-than vehicle-treated PDX (Fig. 4a-b), in line with the reduced cell proliferation observed after elimusertib treatment *in vitro* (Fig. 1). Notably, favorable response to elimusertib treatment, as defined using the RECIST criteria, was associated with low fractions of Ki-67 expressing cells after treatment (overall responding PDX, OR, composed of SD, PR and CR, Fig. 4c). In contrast, in poorly responding PDX, i.e. with progressive disease (PD), differences in Ki-67 staining after elimusertib treatment were not significant (Fig. 4d-i). Similarly, Histone H3 phosphorylation, a marker of mitosis, was lower after elimusertib treatment in 8 out of 9 PDXs classified as responsive (OR, Extended Data fig. 10a-h). Thus, reduced cell proliferation is more pronounced in PDXs responsive to elimusertib. In addition, PDXs were stained for histone variant γH2A.X Ser139 phosphorylation (yH2AX), a marker of DNA damage, and cleaved caspase-3 (Clc3), a marker of apoptosis. In contrast to our *in vitro* results, no significant differences in H2A.X Ser139 phosphorylation or caspase-3 cleavage were observed in PDXs treated with elimusertib compared to vehicle-treated PDXs (Extended Data Fig. 10i-x). This may be because DNA damage induction and apoptosis precede reduced cell proliferation in tumors, hence was not detectable at the end of the treatment period. Thus, reduced Ki-67 expression, indicative of altered tumor cell proliferation, positively correlates with elimusertib response *in vivo* and may serve as a response marker in future clinical trials in which serial biopsies are performed.

**Figure 4.**
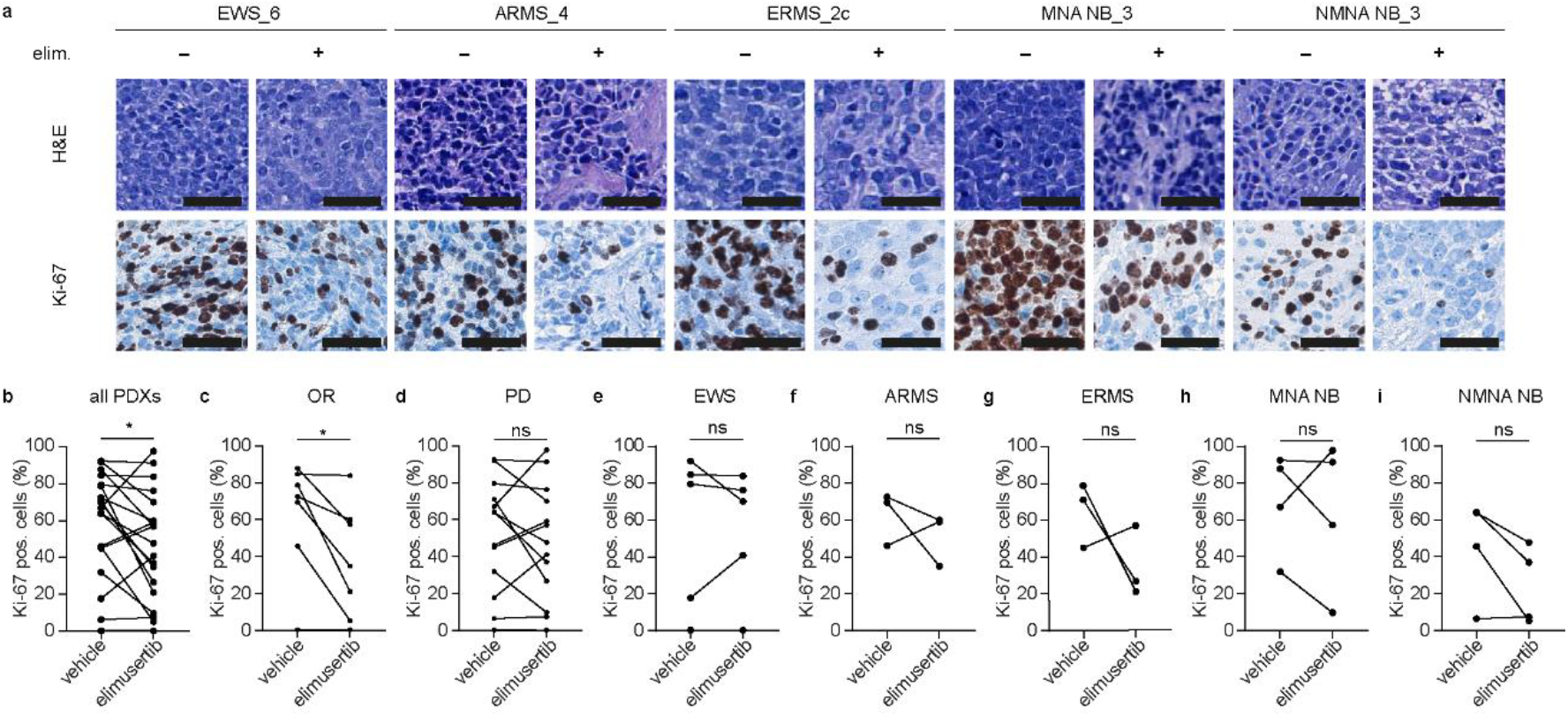
Elimusertib reduces the proliferation rate in PDX models of pediatric solid tumors. **(a)** Exemplary H&E and Ki-67 stainings of EWS, ARMS, ERMS, MNA NB and NMNA NB PDXs treated with elimusertib or vehicle control. **(b-i)** changes in the fraction of Ki-67-expressing cells for all PDXs combined (b), PDXs responding to elimusertib as defined per RECIST (OR, c) and PDXs with progressive disease (PD, d), EWS (e), ARMS (f), ERMS (g), MNA NB (h) and NMNA NB (i). (n = 10; paired, two-sided Student’s t test; error bars represent standard deviation, *P* = 0.0371, 0.0216, 0.4764, 0.9394, 0.4935, 0.2945, 0.7005 and 0.0933 for all PDXs combined, responding PDXs, PDXs with progressive disease, EWS, ARMS, ERMS, MNA NB and NMNA NB, respectively). Scale bar = 40 μm.

### Elimusertib outperforms standard of care treatment in a subset of preclinical pediatric solid tumor models

Pediatric solid tumors are currently treated with a combination of chemotherapeutic agents. In order to evaluate the clinical potential of elimusertib, we aimed to compare the anti-tumor effects of elimusertib in our cohort of PDXs with the effects of current SoC agents. Despite minor differences in exact composition, most pediatric tumors in Europe and the United States are treated in the first line with a combination of topoisomerase inhibitors, mitotic inhibitors, antimetabolites, intercalating and alkylating agents (55-58). The response to the abovementioned chemotherapeutic agents was evaluated using modified RECIST criteria and is also reported in a separate study, in which the detailed molecular features of the PDX used here are presented (Xu et al., currently under consideration elsewhere). We here compared the responses to the SoC chemotherapeutics with the response to elimusertib (Fig. 5a). Notably, most PDXs were relatively unresponsive to SoC chemotherapeutics as monotherapy, which was not associated with prior exposure to these treatments in patients from which PDX were derived. Intriguingly, some of the PDXs that were relatively chemo-resistant responded well to elimusertib, indicating that patients that develop resistance to current SoC treatments may still benefit from elimusertib treatment (Fig. 5). We next compared the changes in PFS following elimusertib treatment to that of SoC chemotherapeutic agents (Fig. 5b-f). Strikingly, elimusertib prolonged the PFS of all ARMS and NMNA NB PDX to a greater extent than any of the SoC agents (Fig. 5). A similarly pronounced prolonged PFS advantage was observed compared to most chemotherapeutic agents tested in ERMS and MNA NB PDX. Only EWS PDX responded similarly to elimusertib as they did to chemotherapy. Thus, our in-depth preclinical response evaluation suggests that elimusertib could have clinically relevant anti-tumor effects in many pediatric tumor entities and may in some cases be superior to currently used treatment options.

**Figure 5.**
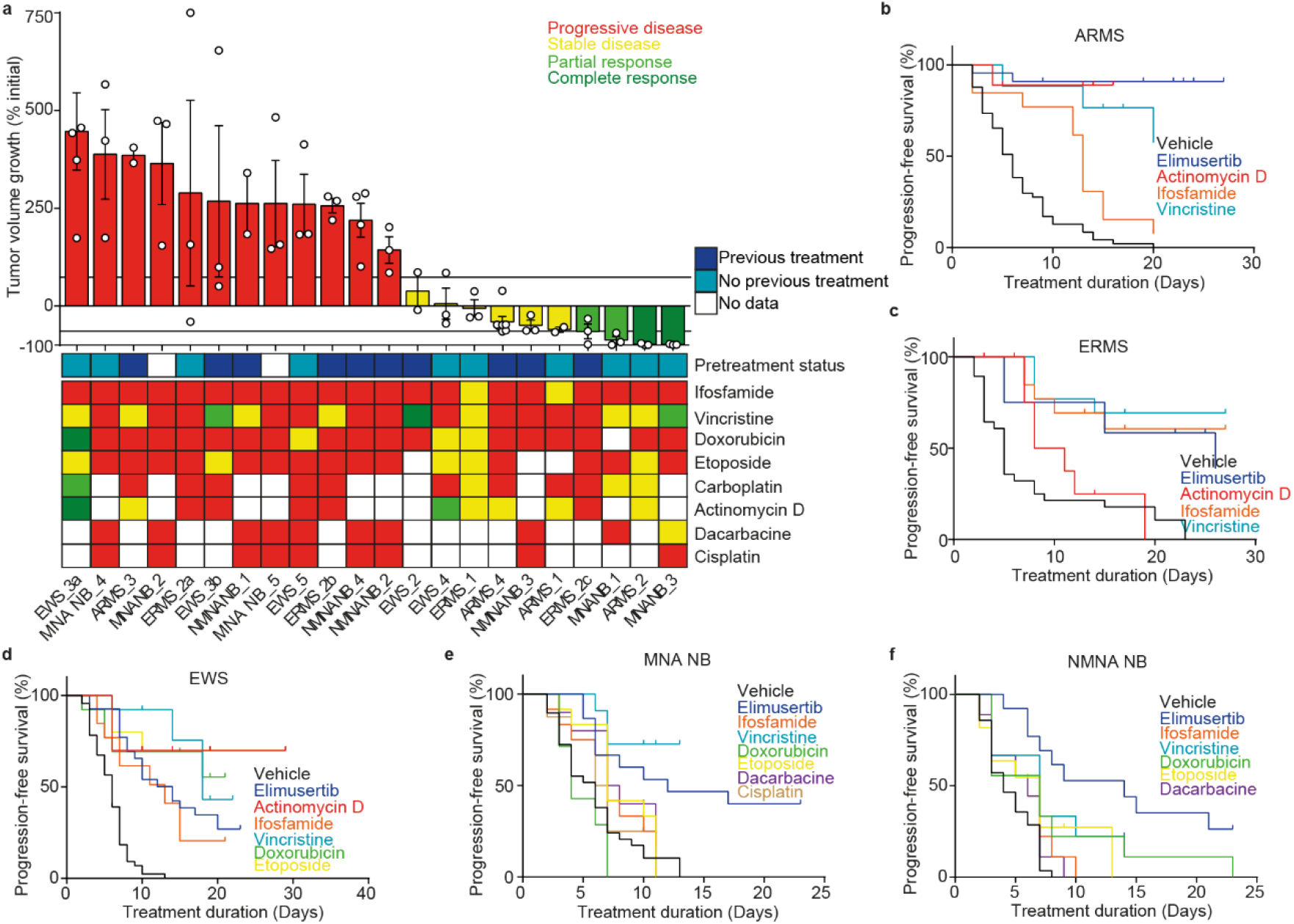
Elimusertib is superior to standard of care treatment in a subset of preclinical pediatric solid tumors models. **(a)** Representation of the tumor volume after elimusertib treatment (top) and response to commonly used chemotherapeutic agents in our cohort of PDX models according to the RECIST criteria in a heatmap (bottom, progressive disease, red; stable disease, yellow; partial response, light green; complete response, dark green;). In dark blue, PDX derived from patients that previously received SoC treatment are marked. (b-f) Kaplan Meier curves comparing the response of tumors to elimusertib, vehicle control treatment, or treatment with standard of care chemotherapeutic agents for ARMS (b), ERMS (c), EWS (d), MNA NB (e), NMNA NB (f).

### Standard of care treatment-associated genomic evolution reveals candidate alterations that render PDXs susceptible to ATR inhibition

As shown *in vitro* (Fig. 1m) and suggested by previous reports (22-28,30), distinct molecular alterations may predict good response to ATR inhibitors. We genetically characterized a subset of the PDX models using whole exome sequencing (results reported in detail in Xu et al., currently under consideration elsewhere). None of the genetic alterations identified in our cohort were associated with therapy response across all or within different entities (Fig. 6a-f). Thus, we focused our analysis on genetic alterations in otherwise near-isogenic PDX pairs derived from the same patients with particularly strong elimusertib response differences (Fig. 6g-h). For example, three ERMS PDX (ERMS_2a-c) derived from subsequent relapses responded very differently to elimusertib, with the best response observed in the PDX derived from the latest relapse (ERMS_2c, Fig. 6b, Extended Data Fig. 3i-k). Intriguingly, mutations in *BRCA1* and *FGFR4* were only detected in the responsive PDX (ERMS_2c) and not in the two PDX derived from earlier clinical timepoints (ERMS_2a+b), suggesting that these mutations occurred later during patient treatment. *BRCA1* deficiency has been implicated in ATR inhibitor response in the past (59,60), suggesting that the improved elimusertib response in the PDX may in part be due to the *de novo BRCA1* mutation. Furthermore ERMS_2b acquired a mutation in *SETD2* during SoC treatment, which has been shown to enhance sensitivity to ATR inhibition in other tumor entitites (30). Additionally, we examined two EWS PDX derived from the same patient (EWS_3a+b). The first model (EWS_3a) was established at diagnosis, whereas the second PDX (EWS_3b) was established from the same patient after neo-adjuvant chemotherapy. Strikingly, the second sample responded better to elimusertib (Fig. 6c, Extended Data Fig. 3n-o), indicating that changes during neo-adjuvant chemotherapy may have enhanced susceptibility to elimusertib. Interestingly, many focal oncogene amplifications (e.g. *MYC, CCND1, MYCN, MDM2*) were detectable in EWS_3b but not EWS_3a (Fig. 6c). In line with previous reports (27,28) and our *in vitro* data (Fig. 1m), *MYCN* was one of the oncogenes mostly amplified in the responsive PDX (Fig. 6a,c). Gene amplifications can arise as a result of genomic instability and can occur in linear or extrachromosomal form (i.e. ecDNA). This raises the possibility that genomic instability and/or the type of gene amplification may influence ATR inhibitor sensitivity.

**Figure 6.**
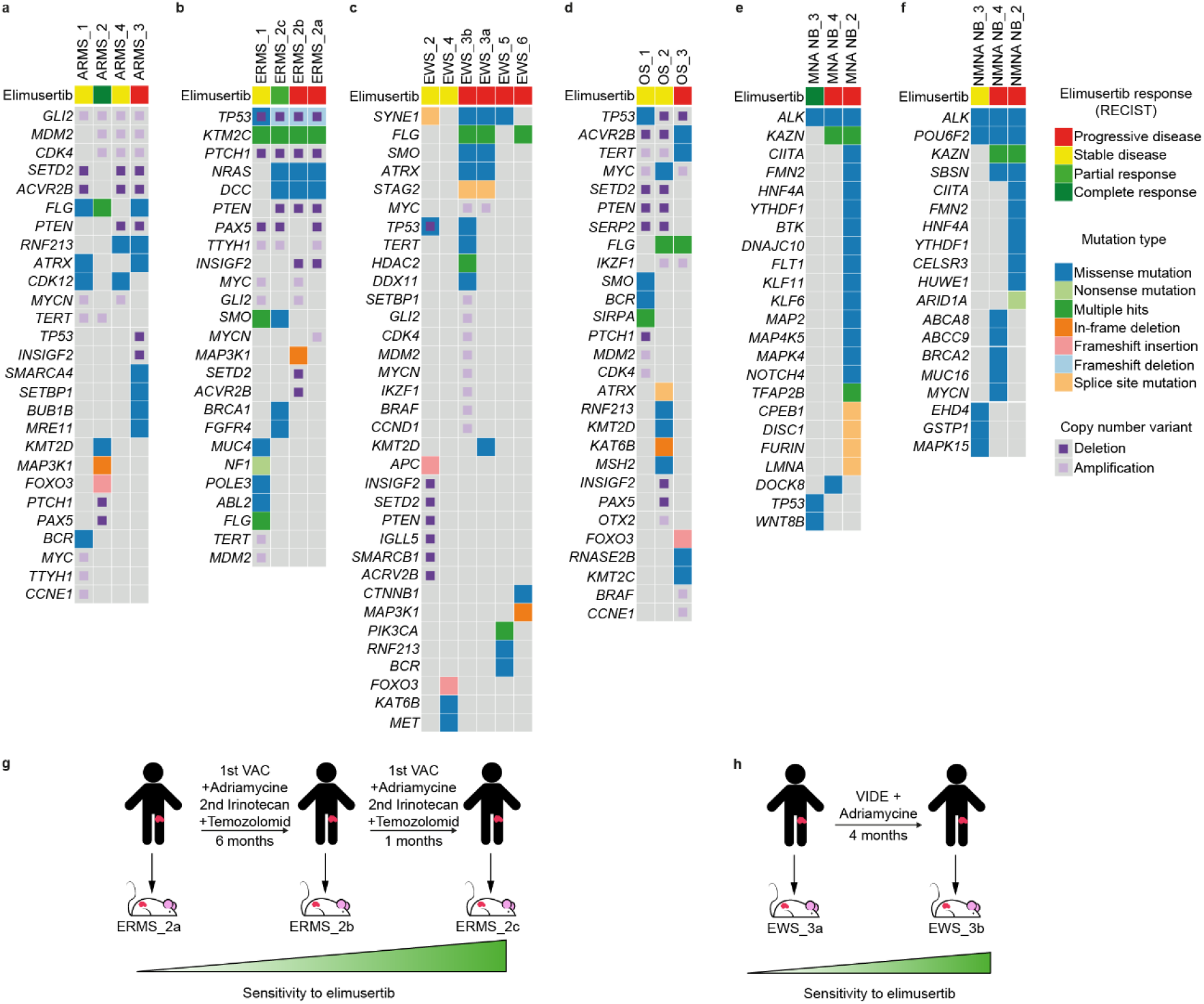
Genomic tumor evolution reveal mutations that are associated with altered response to elimusertib. **(a-f)** Oncoplot showing mutations and CNVs present in PDX models for ARMS (a), ERMS (b), EWS (c), OS (d), MNA NB (e) and NMNA NB (f). **(g)** Timeline and chemotherapy treatment of a patient with ERMS and tumor response to elimusertib of the corresponding PDXs. The first PDX was established from a primary tumor. The patient received a cycle of vincristine, actinomycin D and cyclophosphamide (VAC) complemented with low dosage of doxorubicine. A second line of treatment with irinotecan and temozolomide was added later on. Six months after the first biopsy, a biopsy from a relapsed tumor was used to establish a second PDX, and a new relapse after one month was used for the third PDX. **(h)** Timeline and chemotherapy treatment of a patient with EWS and tumor response to elimusertib of the corresponding PDXs. The first PDX was established from a tumor biopsy used for diagnosis. The patient received a cycle of vincristine, ifosfamide, doxorubicin and etoposide (VIDE) complemented with low dosage of doxorubicine. Four months after the initial biopsy, a biopsy from a relapsed tumor was used to establish a second PDX.

## Discussion

Through an in-depth preclinical assessment of elimusertib’s anti-tumor activity in a broad spectrum of patient-derived pediatric solid tumor models *in vitro* and *in vivo*, we here demonstrate that pharmacological ATR inhibition represents a therapeutic strategy with high clinical potential.

We and others have previously shown that diverse ATR inhibitors exhibit preclinical activity against a subset of ARMS, rhabdoid tumors, OS, EWS, *MYCN*-amplified neuroblastomas and medulloblastomas (24-26,28,29,61), but most of these studies only tested a small number of preclinical models and used ATR inhibitors that are currently not being clinically developed for the use in pediatric patients. In line with our results, the anti-tumor activity of different clinical-stage ATR inhibitors as monotherapy and in combination with other agents has been widely recognized in cancers in adults (21,22,26,38,48,59,62-64).

In contrast to most ATR inhibitors, elimusertib is still in clinical development both for adult and pediatric patients (NCT04095273, NCT04616534, NCT04514497, NCT05071209). Elimusertib’s activity in most pediatric tumor entities, however, has not been assessed comprehensively to date. In an attempt to fill this gap of knowledge, we here performed a preclinical trial using state-of-the art preclinical patient-derived xenografts and broad molecular characterizations, similar to those performed by research consortia like the Pediatric Preclinical Testing Consortium. Compared to previous studies examining the anti-tumor activity ATR inhibitors in small numbers of *in vivo* models, our study provides insights on the inter-tumor response heterogeneity. The response heterogeneity observed in our study mirrors that of many clinical trials for small molecules, suggesting that preclinical trials of this scale may predict clinical responses more closely than preclinical testing using low number of *in vivo* models. High costs of preclinical trials at this scale remain one of the main limitations of such studies. However, we propose that preclinical trials at similar scale as the one performed here should be considered as a standard for preclinical assessments in pediatric oncology.

Previous preclinical trials for various therapeutic interventions conventionally did not compare the effects of the tested intervention to standard of care (SoC) drugs. In fact, very little preclinical data exists for the anti-tumor efficacy of SoC drugs in preclinical patient-derived pediatric tumor models. This is mainly due to the fact that such models were not available to the same extent at the time SoC drugs were first selected for clinical testing. This raises several important questions. Even though many of the same SoC drugs are now considered the clinical gold standard for the treatment of different pediatric patients suffering from molecularly diverse tumor entities, we currently do not know how these SoC drugs perform preclinically. This lack of a true benchmark in preclinical trials creates problems when evaluating the efficacy of new treatment modalities. What anti-tumor effect should we consider as a positive result without such a benchmark? Do we currently set the bar too low or too high for new treatment modalities to be considered successful preclinically? To address these important limitations, we here compared the anti-tumor activity of elimusertib to that of SoC monotherapy in the same PDX models. This revealed that SoC drugs perform surprisingly poor in many PDX when assessing response using clinically relevant read outs and raises the question whether the same drugs would pass the threshold to be approved for clinical testing nowadays. For more details on the molecular profiles and SoC responses of the PDX cohort used here, we refer to a separate publication from our group (Xu et al., currently under consideration elsewhere). We here compared the response to SoC drugs to that of elimusertib, a small molecule inhibitor that very recently entered clinical testing in pediatric patients (NCT05071209). Notably, we observe that elimusertib outperformed most SoC agents in most entities, particularly in ARMS. This is in line with our previous reports describing the exquisite sensitivity of ARMS cells to ATR inhibition, which at least in part seem due to PAX3-FOXO1-induced replication stress (29). We propose that based on both our previous and current studies on ATR inhibitors, patients suffering from ARMS should be designated as a high-priority patient group in which ATR inhibitors should be tested clinically.

Biomarkers predicting clinical response to DDR inhibitors including ATR inhibitors are still scarce. One of the most widely used molecular response predictor used for ATR inhibitors is ATM deficiency (22). Although we cannot exclude that ATM was epigenetically or otherwise compromised, we did not observe an association between the molecular *ATM* status and sensitivity of PDX models to elimusertib (Fig. 6a-f). Our findings stand in line with current clinical trial data showing that a large fraction of patients with ATM deficiency does not respond to ATR inhibitors (33). This suggests that other factors contribute to ATR inhibitor sensitivity. MYCN has been proposed to induce replication stress and sensitize cells to ATR inhibition (26). In line with these reports, *MYCN*-amplified neuroblastoma PDX were amongst the most sensitive to elimusertib. We previously demonstrated that PAX3-FOXO1 expression can sensitize cells to ATR inhibition independent of MYCN expression (29). This raised the question if gene amplification or the type of amplification rather than high oncogene expression may affect ATR inhibitor response. In line with our previous reports, PDX derived from ARMS expressing PAX3-FOXO1, were the most sensitive to elimusertib. Others have reported that fusion oncogene expression in general can sensitize cells ATR inhibition (25,45). In our preclinical trial, however, neither EWS-FLI1-expressing EWS PDX nor CIC-DUX-expressing undifferentiated sarcoma PDX models responded particularly well to elimusertib. The lack of additional CIC-DUX-expressing undifferentiated sarcoma models limits definitive conclusions on the responsiveness of these tumors to elimusertib. As for EWS, we included 8 PDX models in our preclinical trial, 5 of which progressed during elimusertib treatment. This is in stark contrast to the reported sensitivity of EWS cells to ATR inhibition (25,45). We cannot exclude, however, that the previously observed exceptional sensitivity of EWS was specific to the ATR inhibitors tested in these studies and that the chemical or pharmacologic properties of elimusertib influence its activity on EWS cells. Thus, we here provide evidence that ARMS and MYCN-amplified neuroblastomas are most sensitive to elimusertib both *in vitro* and *in vivo*, suggesting patients suffering from these tumor entities may profit from elimusertib treatment.

In summary, elimusertib is active against preclinical patient-derived pediatric solid tumor models. This data supports the initiation of clinical trials with elimusertib in patients with *MYCN*-amplified neuroblastomas and ARMS, and also provides evidence that some tumor entities may not respond as well to elimusertib as previously expected.

## Acknowledgments

We would like to express our gratitude to the patients and families for providing tumor samples to the PDX biobank of Charite – Universitätsmedizin Berlin. We thank Experimental Pharmacology & Oncology GmbH and iPATH.Berlin for their technical support. We want to thank Bayer for providing elimusertib and their financial support for conducting preclinical studies using that drug.

## Extended Data

**Extended Data Figure 1**. Elimusertib represses cell cycle checkpoint activation and induces genomic instability.

**Extended Data Figure 2**. R-loop abundance does not correlate with therapy response to elimusertib in combined pediatric cancer cell lines.

**Extended Data Figure 3**. A cohort of pediatric solid tumor PDXs respond to elimusertib *in vivo*.

**Extended Data Figure 4**. Elimusertib treatment shows limited to no toxicity with regards to body weight development.

**Extended Data Figure 5**. Elimusertib treatment prolongs the overall survival of mice carrying pediatric solid tumors.

**Extended Data Figure 6**. Immunhistochemistry stainings of Ewing Sarcomas.

**Extended Data Figure 7**. Immunhistochemistry stainings of OS, ARMS and ERMS.

**Extended Data Figure 8**. Immunhistochemistry stainings of MNA and NMNA NB.

**Extended Data Figure 9**. Expression patterns of cell cycle, DNA damage and apoptosis markers change with elimusertib treatment.

**Extended Data Figure 10**. Ki-67 expression is reduced upon treatment with elimusertib in responding PDXs.

**Extended Data Figure 11**. Baseline IHC markers for proliferation, DNA damage and apoptosis show no correlation with relative tumor volume after treatment.

**Extended Data Table 1**. IC50 and AUC values of all used pediatric cancer cell lines.

**Extended Data Table 2**. PDX characterization with regards to tumor status, biopsy location, metastases, grading/staging, age and sex of the patients.

**Extended Data Table 3**. Statistical data for all tumor volume curves displayed in Extended Data Figure 3.

**Exteded Data Table 4**. Quantifications of IHC markers in RMS.

**Extended Data Table 5**. Quantifications of IHC markers in EWS and OS_1.

**Extended Data Table 6**. Quantifications of IHC markers in NB.

