## Extended Data for "Elimusertib outperforms standard of care chemotherapy in preclinical patient-derived pediatric solid tumor models"

[7]iPATH.Berlin – Core Unit Immunopathology for Experimental Models, Charité Berlin, corporate member of Freie Universität Berlin, Humboldt-Universität zu Berlin, Berlin, Germany

[8]Department of Pediatrics, Memorial Sloan Kettering Cancer Center, New York City, NY, USA.

[9]Bayer AG, Berlin, Germany.

[10]Berlin Institute of Health, 10178 Berlin, Germany.

[11]German Cancer Consortium (DKTK), partner site Berlin, and German Cancer Research Center (DKFZ), Heidelberg, Germany.

[#]These authors contributed equally.

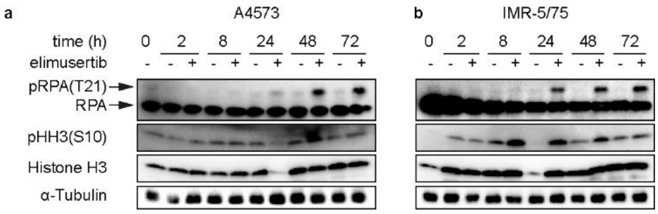


**Extended Data Figure 1. Elimusertib represses cell cycle checkpoint activation and induces genomic instability. (a-b)** Western Immunoblotting of RPA32, pRPA(T21), Histone H3 and pHH3(S10) in A4573 (a) and IMR-5/75 cells (b) treated over time with elimusertib (30 nM). RPA: replication protein A 32; pRPA(T21): replication protein A 32, phosphorlyated in position 21 (threonine); pHH3(S10): Histone H3 phosphorylated in position 10 (serine).

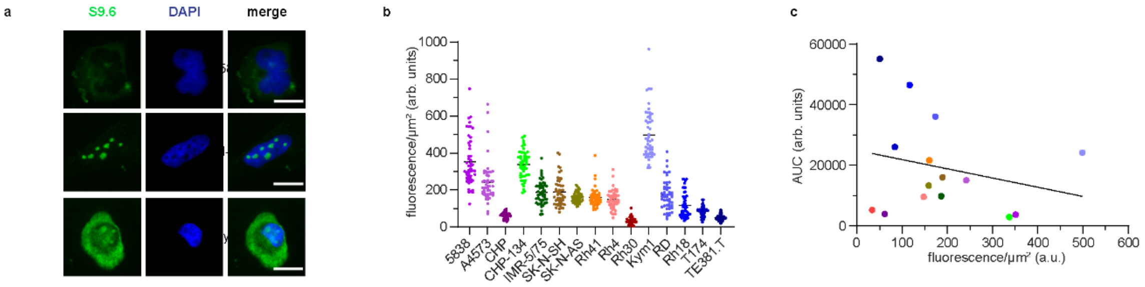


**Extended Data Figure 2. R-loop abundance does not correlate with therapy response to elimusertib in combined pediatric cancer cell lines. (a)** Representative immunofluorescence microphotographs of R-loops (S9.6) and DAPI signal in cells. **(b)** Quantification of R-loop fluorescence in cell lines **(c)** Correlation between AUC for elimusertib and R-loop signal in cell lines ($R^2 = 0.05853$, $p = 0.8020$). arb. units: arbitrary units. AUC: area under the curve. S9.6: monoclonal antibody targeting DNA-RNA hybrids (R-loops), clone S9.6. Scale bar = 20 µm.

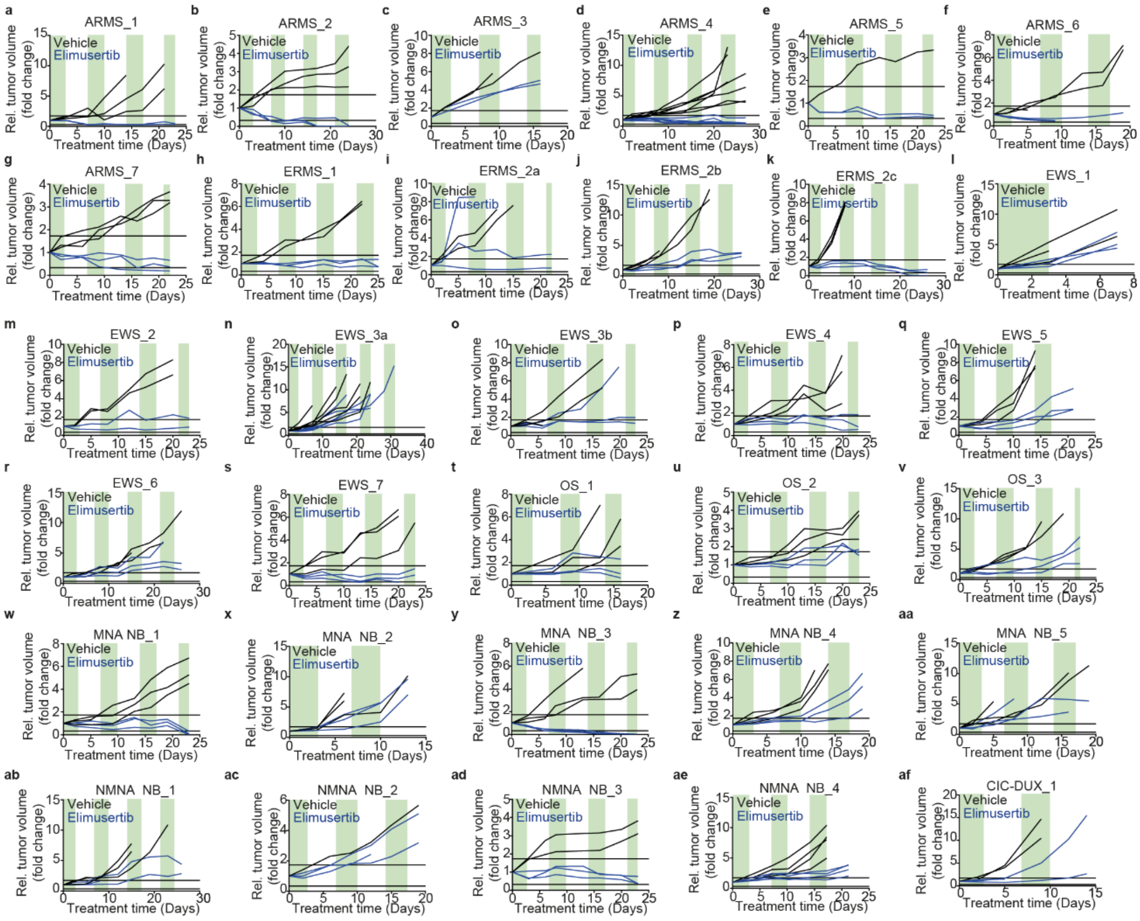


**Extended Data Figure 3. A cohort of pediatric solid tumor PDXs respond to elimusertib *in vivo*. (a-af)** Relative tumor volumes over the course of elimusertib treatment in 7 ARMS (a-g), 4 ERMS (h-k), 8 EWS (l-s), 3 OS (t-v), 5 MNA NB (w-aa), 4 NMNA NB (ab-ae) and one CIC-DUX model (af).

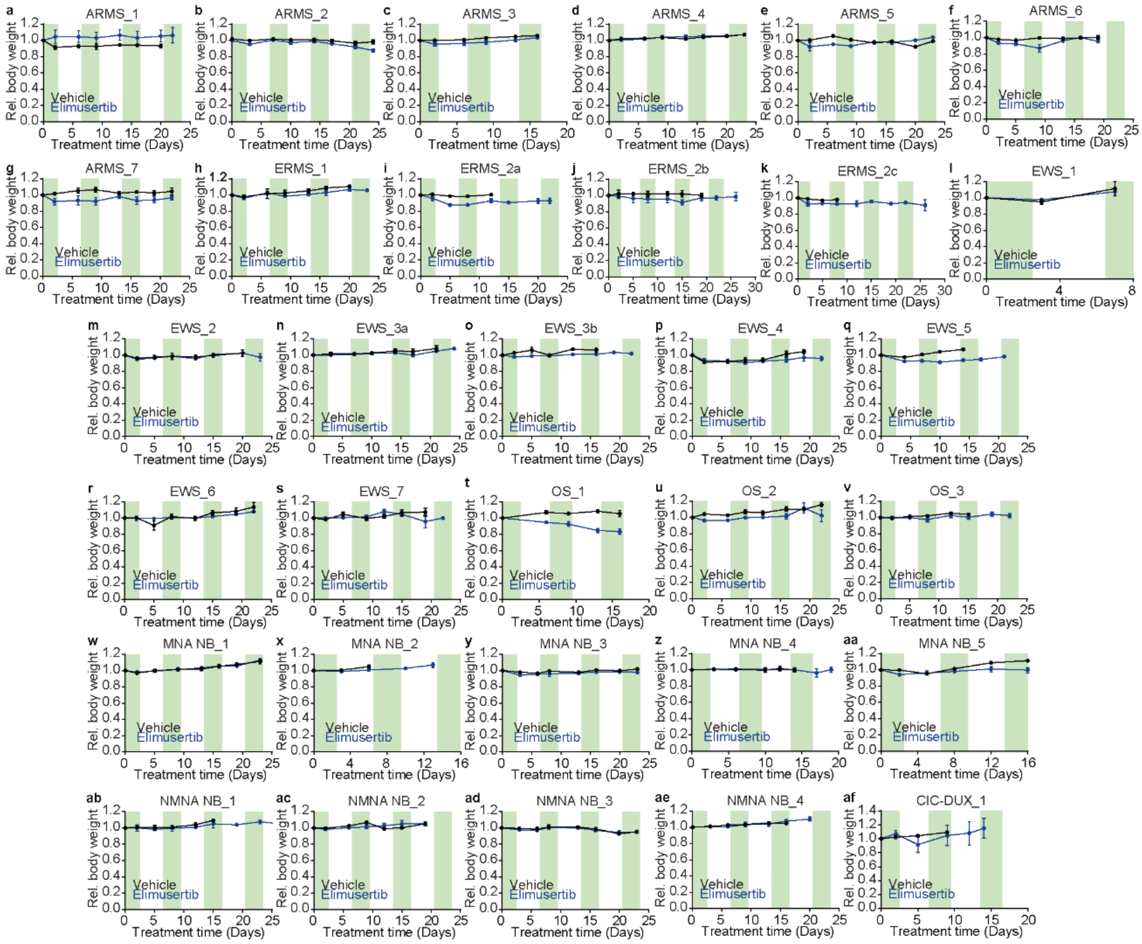


**Extended Data Figure 4. Elimusertib treatment shows limited to no toxicity with regards to body weight development. (a-af)** body weight curves of all ARMS (a-g), ERMS (h-k), EWS (l-s), OS (t-v), MNA NB (w-aa), NMNA NB (ab-ae) and one CIC-DUX model (af).

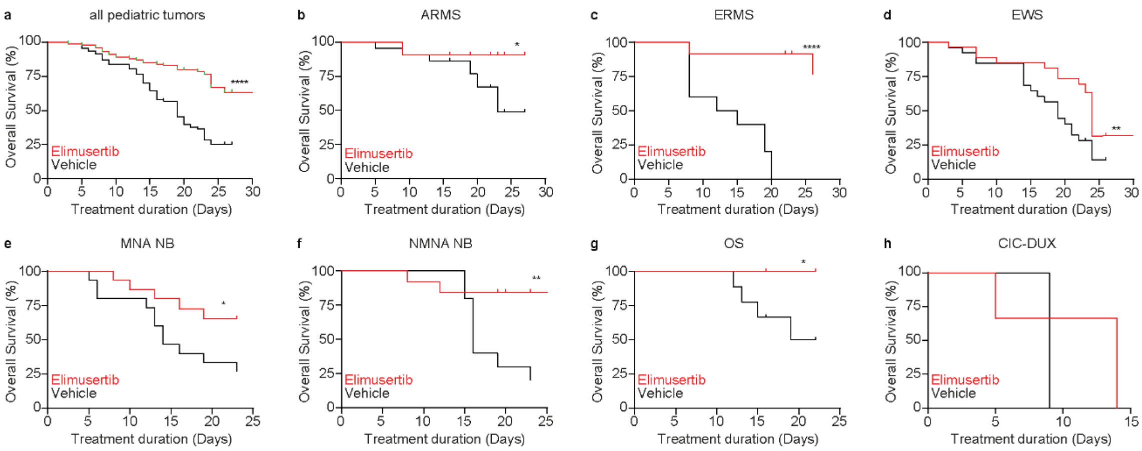


**Extended Data Figure 5. Elimusertib treatment prolongs the overall survival of mice carrying pediatric solid tumors. (a-h)** Kaplan Meier curve showing the overall survival of mice receiving elimusertib across tumor types (a, $n_{total}$ = 195, $P$ = < 0.0001), ARMS (b, $n_{total}$ = 44, $P$ = 0.0121), ERMS (c, $n_{total}$ = 22, $P$ = < 0.0001), EWS (d, $n_{total}$ = 53, $P$ = 0.0086), MNA NB (e, $n_{total}$ = 30, $P$ = 0.0447), NMNA NB (f, $n_{total}$ = 23, $P$ = 0.0099), OS (g, $n_{total}$ = 18, $P$ = 0.0272) and CIC-DUX sarcoma (h, $n_{total}$ = 5, $P$ = 0.4281). Log-rank tests were performed for statistical comparison.

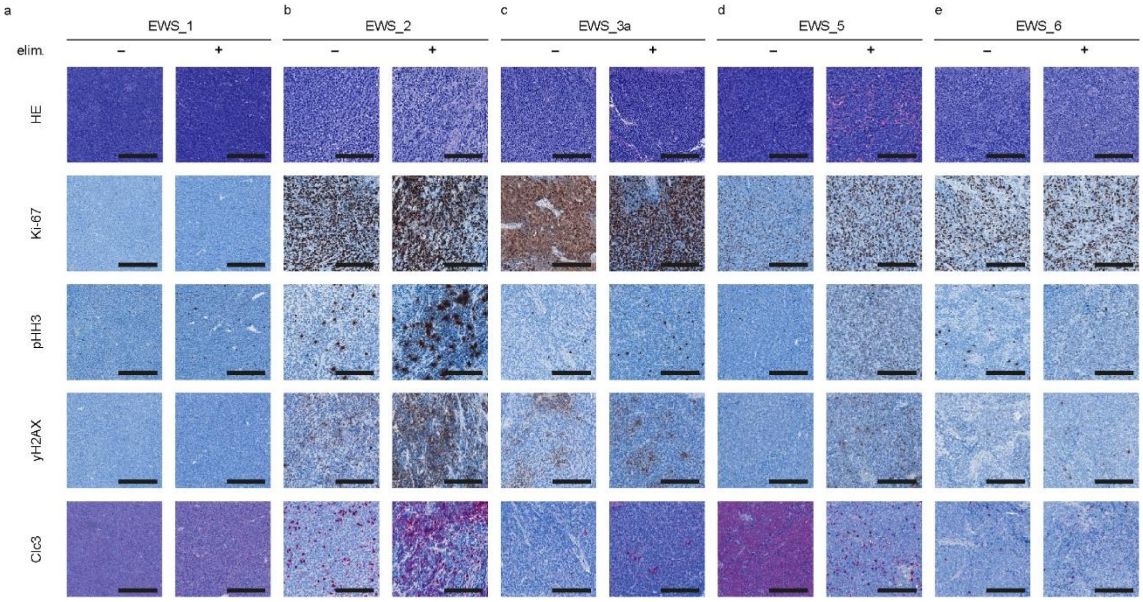


**Extended Data Figure 6. Immunohistochemistry stainings of Ewing Sarcomas. (a-e)** Haematoxylin and eosin (HE) stainings as well as immunohistochemistry stainings of Ki-67, phospho-Histone H3 (S10) (pHH3), phospho-H2A.X (S139) (yH2AX) and cleaved caspase-3 (Clc3) in PDX-models EWS-1 (a), EWS-2 (b), EWS-3a (c), EWS-5 (d) and EWS-6 (e). Scale bar = 200 µm.

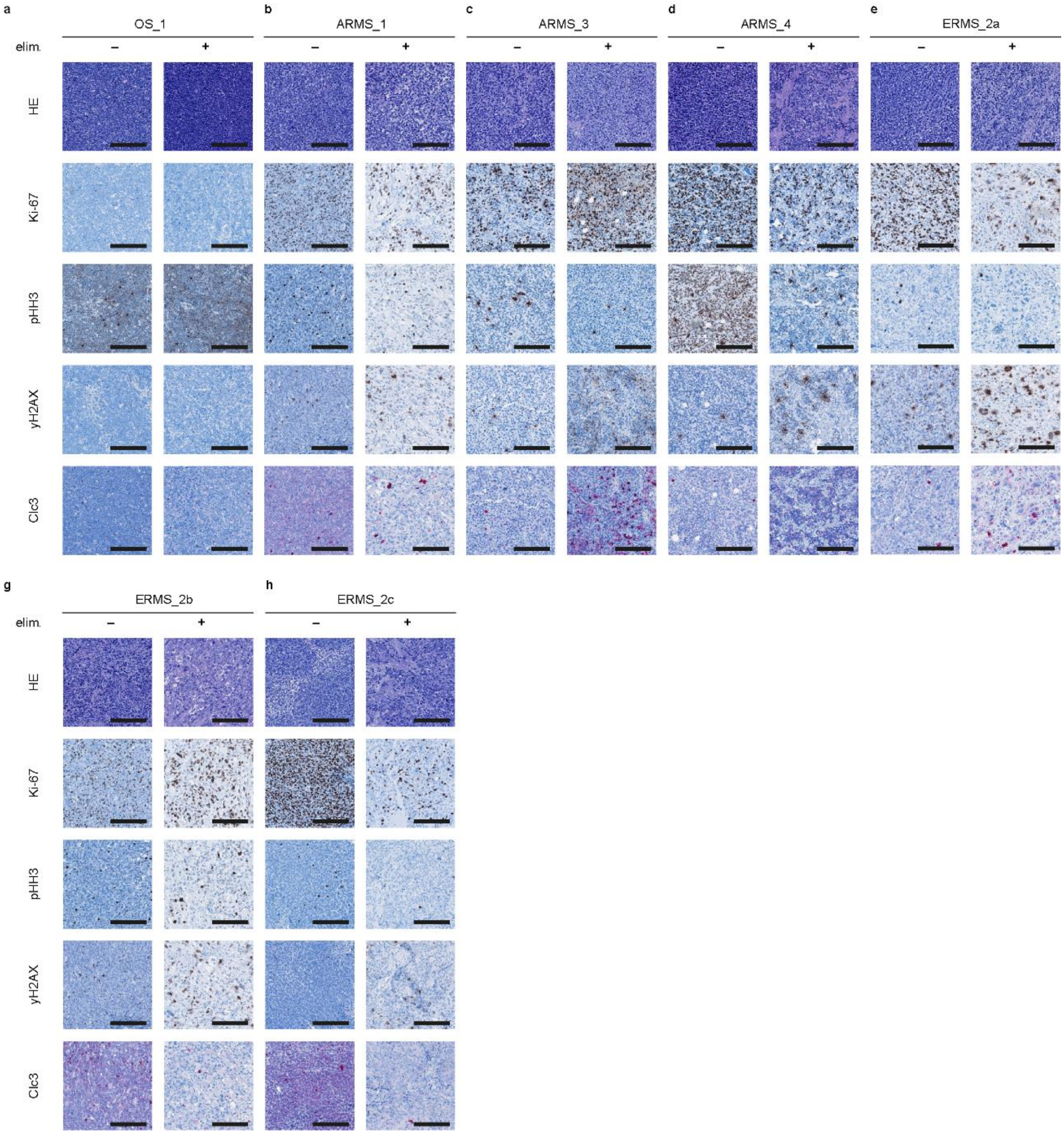


**Extended Data Figure 7. Immunohistochemistry stainings of OS, ARMS and ERMS. (a-h)** Haematoxylin and eosin (HE) stainings as well as IHC stainings of Ki-67, phospho-Histone H3 (S10) (pHH3), phospho-H2A.X (S139) (yH2AX) and cleaved caspase-3 (Clc3) in PDX-models OS-1 (a), ARMS-1 (b), ARMS-2 (c), ARMS-3 (d), ARMS-4 (e), ERMS-2a (f), ERMS-2b (g) and ERMS-2c (h). Scale bar = 200 µm.

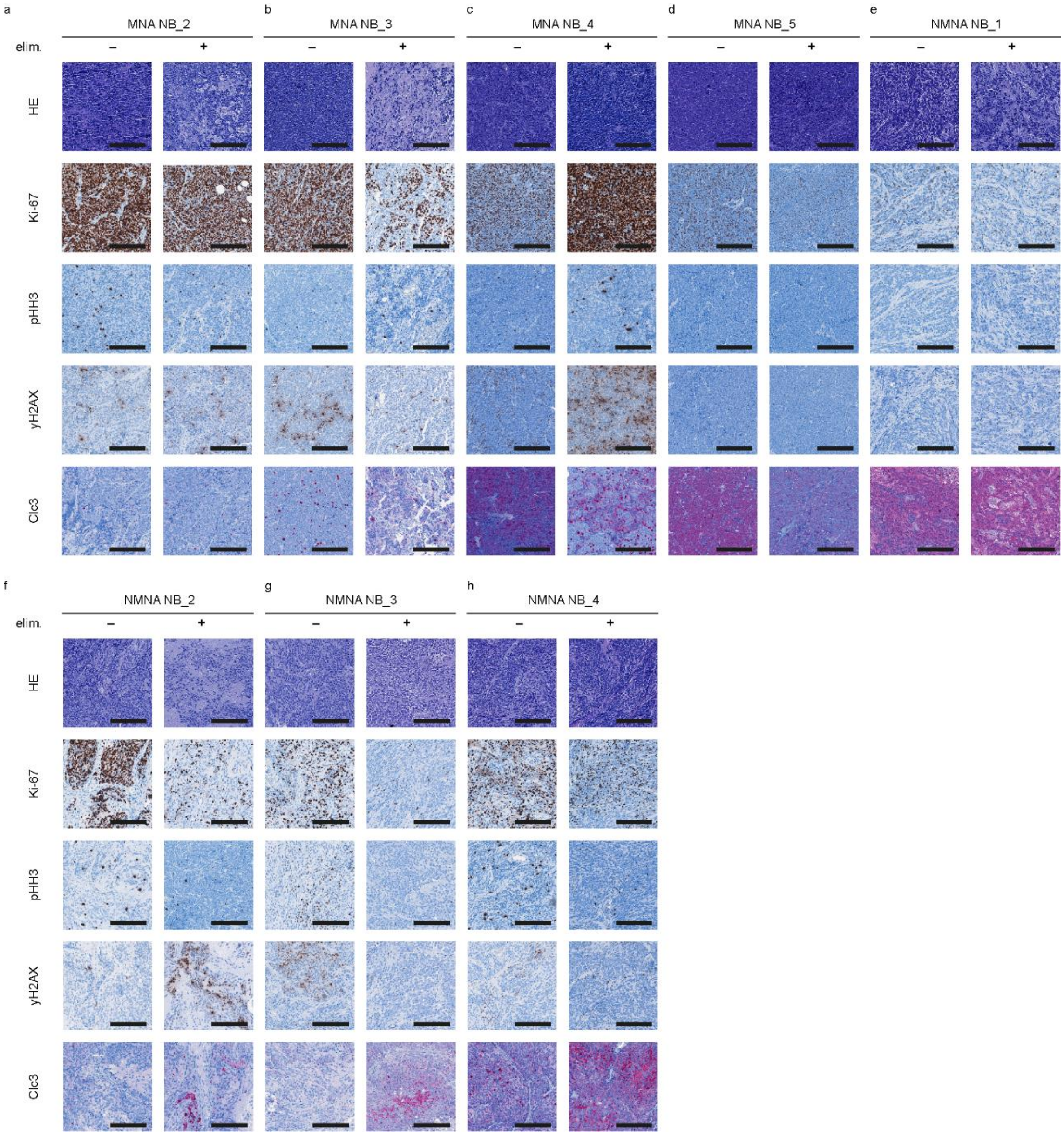


**Extended Data Figure 8. Immunohistochemistry stainings of MNA and NMNA NB. (a-h)** Haematoxylin and eosin (HE) stainings as well as IHC stainings of Ki-67, phospho-Histone H3 (S10) (pHH3), phospho-H2A.X (S139) (yH2AX) and cleaved caspase-3 (Clc3) in PDX-models MNA NB-2 (a), MNA NB-3 (b), MNA NB-4 (c), MNA NB-5 (d), NMNA NB-1 (e), NMNA NB-2 (f), NMNA NB-3 (g) and NMNA NB-4 (h). Scale bar = 200 µm.

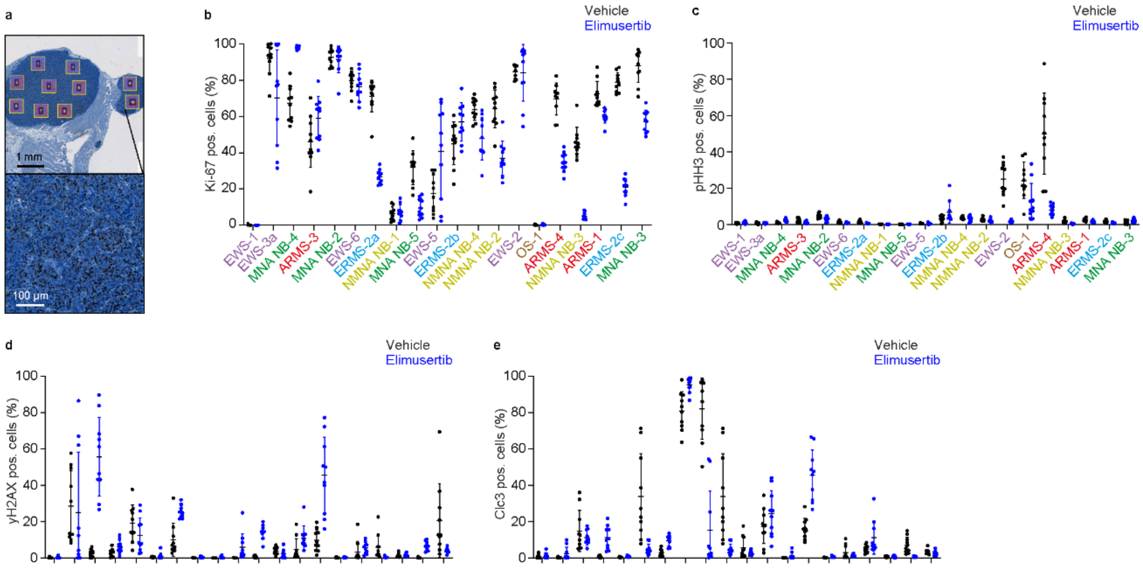


**Extended Data Figure 9. Expression patterns of cell cycle, DNA damage and apoptosis markers change with elimusertib treatment. (a)** Exemplary quantification of Ki-67 in an IHC staining of a PDX. **(b-d)** Dot plot of Ki-67 (b), phosho-histone H3 (S10) (c), phospho-H2AX (S139) (d) and cleaved caspase-3 quantifications (e). Mean, SD and p values are listed in Extended Data Table 4-6.

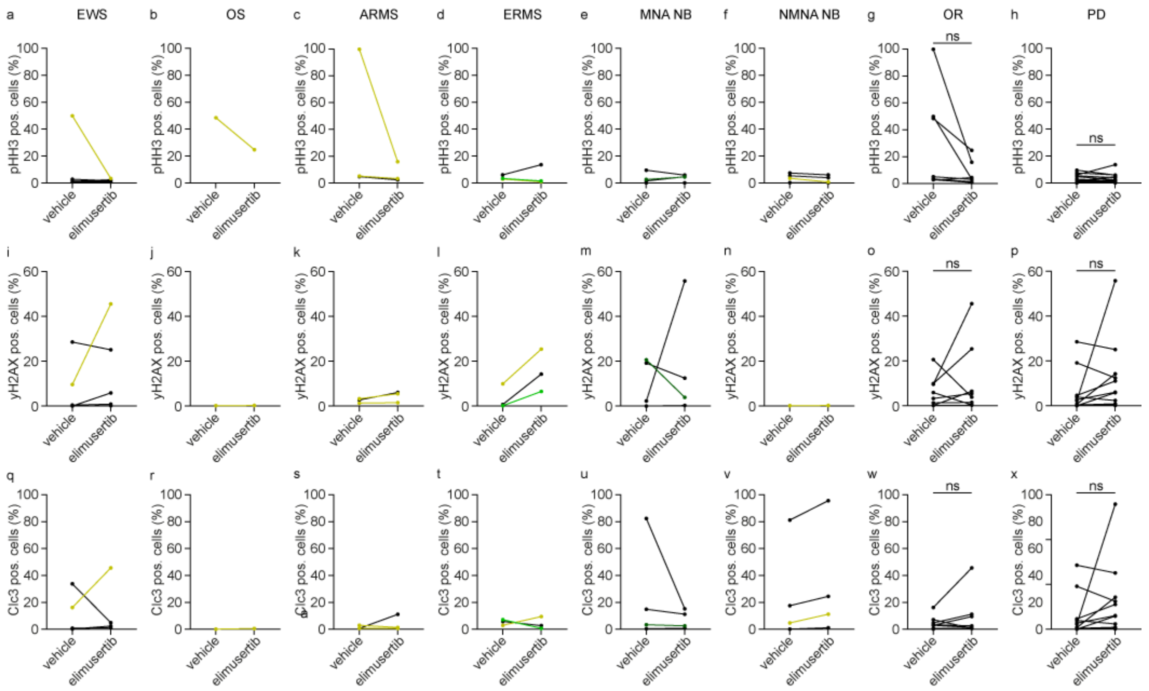


**Extended Data Figure 10. Ki-67 expression is reduced upon treatment with elimusertib in responding PDXs. (a-f)** Changes in the mean expression of phospho-Histone H3(S10) upon treatment with elimusertib in EWS (a), OS (b), ARMS (c), ERMS (d), MNA NB (e) and NMNA NB (f). **(g-h)** Changes in the mean expression of phospho-Histone H3(S10) in overall responding PDXs (g) and PDXs showing a progressive disease under treatment (h). **(i-n)** Changes in the mean expression of phospho-H2AX(S139) upon treatment with elimusertib in EWS (i), Osteosarcoma (j), ARMS (k), ERMS (l), MNA NB (m) and NMNA NB (n). **(o-p)** Changes in the mean expression phospho-H2AX(S139) in overall responding PDXs (o) and PDXs showing a progressive disease under treatment (p). **(q-v)** Changes in the mean expression of cleaved caspase-3 upon treatment with elimusertib in EWS (q), Osteosarcoma (r), ARMS (s), ERMS (t), MNA NB (u) and NMNA NB (v). **(w-x)** Changes in the mean expression of cleaved caspase-3 in overall responding PDXs (w) and PDXs showing a progressive disease under treatment (x). Two-sided, paired t-tests were performed for statistical analysis.

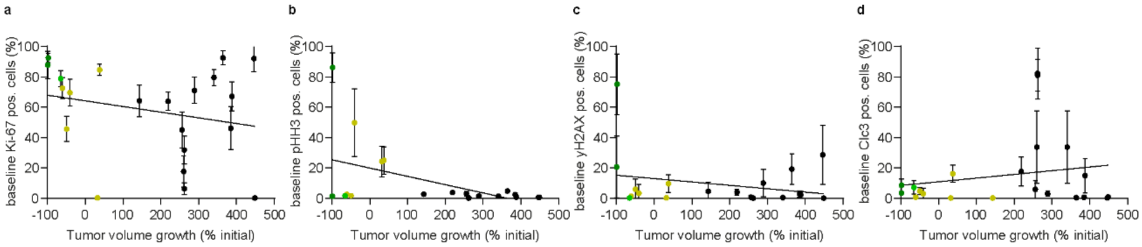


**Extended Data Figure 11. Baseline IHC markers for proliferation, DNA damage and apoptosis show no correlation with relative tumor volume after treatment. (a-d)** Correlation between the baseline Ki-67 (a), pHH3 (b), yH2AX (c) and Clc3 levels (d) and the relative tumor volume after treatment with elimusertib. $R^2$ = 0.05522, 0.2472, 0.06322 and 0.03729 for Ki-67, pHH3, yH2AX and Clc3 respectively. *P* = 0.3053, 0.0218, 0.2716 and 0.7360 for Ki-67, pHH3, yH2AX and Clc3 respectively.

**Extended Data Table 1. IC50 and AUC values of all used pediatric cancer cell lines.**

| | Cell line | IC50 [nM] | AUC (a.u.) |
|---|---|---|---|
| *MYCN* amplified Neuroblastoma | BE(2)-C | 139.4 | 31353 |
| | CHP-134 | 10.5 | 2854 |
| | CHP-212 | 41.96 | 29209 |
| | IMR5/75 | 18.76 | 9798 |
| | Kelly | 11.79 | 2365 |
| | LAN-5 | 14.58 | 8995 |
| | NB-01 | 97.1 | 30679 |
| | NB-69 | 6.411 | 2988 |
| | NGP | 5.666 | 6521 |
| | SK-N-BE | 33.95 | 21999 |
| | SK-N-DZ | 14.05 | 7985 |
| | TR-14 | 84.81 | 24348 |
| Non-*MYCN* amplified Neuroblastoma | GI-ME-N | 61.72 | 34441 |
| | LAN-6 | 153.6 | 37730 |
| | SH-EP | 67.38 | 27639 |
| | SH-SY5Y | 2.315 | 4763 |
| | SK-N-AS | 26.02 | 13294 |
| | SK-N-FI | 19.85 | 17089 |
| | SK-N-SH | 60.22 | 16003 |
| Ewing Sarcoma | 5838 | 13.52 | 3889 |
| | A4573 | 38.73 | 15037 |
| | CADO-ES1 | 14.12 | 10481 |
| | CHP | 13.93 | 3680 |
| | JR | 253.2 | 44576 |
| | SB | 45.7 | 20423 |
| | SK-N-MC | 5.597 | 8143 |
| | TC-71 | 18.96 | 5775 |
| Alveolar rhabdomyosarcoma | KFR | 64.44 | 25973 |
| | Rh4 | 45.19 | 9559 |
| | Rh5 | 245.7 | 42187 |
| | Rh28 | 5.247 | 17675 |
| | Rh30 | 12.31 | 5242 |
| | Rh41 | 69.36 | 21602 |
| | RMS | 22.49 | 16545 |
| Embryonal rhabdomyosarcoma | Kym1 | 69.01 | 24184 |
| | RD | 142.5 | 36058 |
| | Rh18 | 302.6 | 46485 |
| | T174 | 62.24 | 19897 |
| | TE381.T | 450.8 | 55101 |
| Non-cancer cells | RPE | 132.9 | 34051 |
| | BJ | 213.4 | 43900 |

**Extended Data Table 2. PDX characterization with regards to tumor status, biopsy location, metastases, grading/staging, age and sex of the patients.** L: lymph node metastasis; M: distant metastasis; L+M: lymph node and distant metastasis. Adr. Gland: adrenal gland. N/A: not applicable/no information; Y: age in years; Mo: Age in months;

| PDX name | Tumor Status | Biopsy Location | Metastases | Grading/Staging | Age (Y/Mo) | Sex |
|---|---|---|---|---|---|---|
| ARMS_1 | primary | foot | L+M | very high risk | 12 Y | male |
| ARMS_2 | primary | hand | - | high risk | 10 Y | female |
| ARMS_3 | 1st relapse | pararectal | L+M | very high risk | 15 Y | female |
| ARMS_4 | 1st relapse | foot | L+M | very high risk | 16 Y | female |
| ARMS_5 | relapse | M. abducens | N/A | N/A | 10 Y | female |
| ARMS_6 | relapse | bladder | N/A | N/A | 3 Y | female |
| ARMS_7 | 1st relapse | head | L+M | N/A | 6 Y | N/A |
| ERMS_1 | primary | pelvis | M | very high risk | 12 Y | male |
| ERMS_2a | primary | M. masseter | - | standard risk | 5 Mo | female |
| ERMS_2b | 1st relapse | - | - | N/A | 11 Mo | female |
| ERMS_2c | 2nd relapse | - | - | N/A | 1 Y | female |
| EWS_1 | primary | thorax – soft tissue | - | N/A | 3 Y | female |
| EWS_2 | 1st local recurrence | pelvis | M | N/A | 16 Y | female |
| EWS_3a | primary | pelvis | - | N/A | 5 Y | male |
| EWS_3b | primary (post treatment) | pelvis | - | N/A | 6 Y | male |
| EWS_4 | primary | thorax | L+M | N/A | 11 Y | female |
| EWS_5 | primary | lower leg – soft tissue | - | N/A | 8 Y | male |
| EWS_6 | primary | femur | L | high-grade | 17 Y | male |
| EWS_7 | N/A | N/A | N/A | N/A | N/A | N/A |
| MNA NB_1 | primary | adr.gland | N/A | III | 1 Y | female |
| MNA NB_2 | N/A | N/A | N/A | N/A | N/A | N/A |
| MNA NB_3 | primary | adr. gland | N/A | IV | 1 Y | male |
| MNA NB_4 | primary | adr. gland | N/A | IV | 7 Y | male |
| MNA NB_5 | N/A | N/A | N/A | N/A | N/A | N/A |
| NMNA NB_1 | relapse | thorax | N/A | IV | 14 Y | male |
| NMNA NB_2 | relapse | abdomen | N/A | IV | 2 Y | female |
| NMNA NB_3 | relapse | abdomen | N/A | III | 10 Mo | male |
| NMNA NB_4 | relapse | N/A | N/A | N/A | 6 Y | male |
| OS_1 | primary | pelvis | - | high-grade | 7 Y | female |
| OS_2 | primary | humerus | L | high-grade | 11 Y | female |
| OS_3 | primary | femur | - | high-grade | 16 Y | female |
| CIC_DUX | primary | gluteus | M | high-grade | 9 Y | male |

**Extended Data Table 3. Statistical data for all tumor volume curves displayed in Figure 2b and Extended Data Figure 2.** TV: tumor volume.

| PDX model | $n_{vehicle}$ | $n_{elimusertib}$ | p value | $TV_{vehicle}$ (fold change) | $TV_{elimusertib}$ (fold change) |
|---|---|---|---|---|---|
| ARMS_1 | 3 | 2 | 0,121603 | 5,132 | 0,3087 |
| ARMS_2 | 3 | 3 | 0,007189 | 3,279 | 0,01889 |
| ARMS_3 | 2 | 2 | 0,097023 | 5,248 | 3,557 |
| ARMS_4 | 7 | 7 | 0,000331 | 7,481 | 0,6123 |
| ARMS_5 | 1 | 2 | 0,015470 | 3,336 | 0,4214 |
| ARMS_6 | 3 | 3 | 0,003844 | 1,652 | 0,5782 |
| ARMS_7 | 3 | 3 | 0,000118 | 3,381 | 0,4054 |
| ERMS_1 | 2 | 3 | 0,000275 | 6,338 | 1,190 |
| ERMS_2a | 2 | 3 | 0,990833 | 3,849 | 3,888 |
| ERMS_2b | 3 | 3 | 0,006182 | 3,530 | 1,443 |
| ERMS_2c | 3 | 3 | 0,000007 | 7,959 | 1,334 |
| EWS_1 | 2 | 3 | 0,136776 | 3,819 | 1,943 |
| EWS_2 | 2 | 2 | 0,031454 | 7,465 | 1,576 |
| EWS_3a | 3 | 4 | 0,049562 | 3,173 | 1,591 |
| EWS_3b | 2 | 2 | 0,258522 | 1,786 | 1,134 |
| EWS_4 | 3 | 3 | 0,040344 | 5,170 | 1,249 |
| EWS_5 | 3 | 3 | 0,001135 | 8,111 | 2,015 |
| EWS_6 | 2 | 3 | 0,116805 | 4,796 | 3,090 |
| EWS_7 | 3 | 3 | 0,014710 | 5,284 | 0,6298 |
| MNA NB_1 | 3 | 3 | 0,001288 | 5,499 | 0,1315 |
| MNA NB_2 | 3 | 2 | 0,079440 | 5,680 | 2,692 |
| MNA NB_3 | 2 | 3 | 0,013872 | 4,031 | 0,2714 |
| MNA NB_4 | 2 | 3 | 0,010926 | 5,695 | 2,090 |
| MNA NB_5 | 2 | 2 | 0,426988 | 3,380 | 2,359 |
| NMNA NB_1 | 3 | 2 | 0,252488 | 6,008 | 3,363 |
| NMNA NB_2 | 1 | 3 | 0,359519 | 3,225 | 2,432 |
| NMNA NB_3 | 2 | 3 | 0,002491 | 3,464 | 0,5056 |
| NMNA NB_4 | 4 | 4 | 0,002580 | 7,959 | 2,198 |
| OS_1 | 3 | 3 | 0,267491 | 3,834 | 1,691 |
| OS_2 | 3 | 3 | 0,026347 | 3,374 | 1,682 |
| OS_3 | 3 | 3 | 0,018217 | 5,138 | 2,134 |
| CIC-DUX | 2 | 3 | 0,005353 | 4,269 | 1,314 |

**Extended Data Table 4. Quantifications of IHC markers in RMS.**

| | | PDX model | ARMS_1 | ARMS_2 | ARMS_3 | ARMS_4 | ERMS_2a | ERMS_2b | ERMS_2c |
|---|---|---|---|---|---|---|---|---|---|
| Ki-67 | veh. | mean (%) | 72.522 | 92.558 | 46.150 | 69.591 | 71.101 | 45.012 | 78.837 |
| | | SD (%) | 6.562 | 2.817 | 13.467 | 8.310 | 8.263 | 11.187 | 5.026 |
| | eli. | mean (%) | 59.984 | - | 59.092 | 34.877 | 26.694 | 56.963 | 20.921 |
| | | SD (%) | 3.471 | - | 11.358 | 5.041 | 3.605 | 10.014 | 4.609 |
| | | p value | 0.0007 | - | 0.0101 | 9E-08 | 1E-08 | 0.0617 | 4E-11 |
| pHH3 | veh. | mean (%) | 2.571 | 86.205 | 2.350 | 49.892 | 1.419 | 2.988 | 1.666 |
| | | SD (%) | 0.696 | 9.146 | 1.067 | 21.176 | 0.616 | 1.622 | 0.542 |
| | eli. | mean (%) | 1.597 | - | 1.147 | 7.943 | 0.444 | 6.777 | 0.718 |
| | | SD (%) | 0.484 | - | 0.621 | 2.744 | 0.223 | 5.659 | 0.289 |
| | | p value | 0.0111 | - | 0.0108 | 0.0002 | 0.0019 | 0.1294 | 0.0035 |
| yH2AX | veh. | mean (%) | 1.326 | 75.223 | 2.715 | 3.278 | 9.957 | 0.762 | 0.159 |
| | | SD (%) | 1.000 | 19.023 | 2.112 | 5.558 | 8.651 | 0.472 | 0.207 |
| | eli. | mean (%) | 1.521 | - | 6.088 | 5.510 | 25.392 | 14.273 | 6.514 |
| | | SD (%) | 0.792 | - | 3.462 | 2.975 | 3.275 | 3.618 | 2.337 |
| | | p value | 0.6706 | - | 0.0217 | 0.2889 | 0.0011 | 2E-06 | 2E-05 |
| Clc3 | veh. | mean (%) | 0.681 | 8.550 | 0.665 | 2.976 | 2.980 | 5.765 | 7.117 |
| | | SD (%) | 0.316 | 4.009 | 0.464 | 3.622 | 1.681 | 5.582 | 4.227 |
| | eli. | mean (%) | 0.748 | - | 11.086 | 1.367 | 9.480 | 2.723 | 0.672 |
| | | SD (%) | 0.447 | - | 6.009 | 0.636 | 2.822 | 1.098 | 0.312 |
| | | p value | 0.6906 | - | 0.0006 | 0.2025 | 6E-05 | 0.138 | 0.0016 |

**Extended Data Table 5. Quantifications of IHC markers in EWS and OS_1.**

| | | PDX model | EWS_1 | EWS_2 | EWS_3a | EWS_5 | EWS_6 | OS_1 |
|---|---|---|---|---|---|---|---|---|
| Ki-67 | veh. | mean (%) | 0.149 | 84.707 | 92.125 | 17.632 | 79.648 | 0.174 |
| | | SD (%) | 0.234 | 3.545 | 8.380 | 9.789 | 5.152 | 0.174 |
| | eli. | mean (%) | 0.000 | 84.005 | 70.143 | 40.915 | 76.288 | 0.260 |
| | | SD (%) | 0.000 | 14.793 | 25.213 | 24.512 | 7.156 | 0.306 |
| | | p value | 0.0884 | 0.899 | 0.0234 | 0.0062 | 0.333 | 0.4909 |
| pHH3 | veh. | mean (%) | 0.596 | 4.625 | 0.615 | 0.405 | 1.365 | 24.193 |
| | | SD (%) | 0.371 | 3.095 | 0.387 | 0.179 | 0.480 | 9.485 |
| | eli. | mean (%) | 1.201 | 24.928 | 0.645 | 0.334 | 0.807 | 12.330 |
| | | SD (%) | 0.504 | 8.640 | 0.343 | 0.363 | 0.324 | 9.619 |
| | | p value | 0.0062 | 6E-05 | 0.8595 | 0.645 | 0.0268 | 0.0382 |
| yH2AX | veh. | mean (%) | 0.082 | 9.616 | 28.548 | 0.035 | 0.406 | 0.192 |
| | | SD (%) | 0.245 | 5.736 | 18.542 | 0.076 | 0.215 | 0.156 |
| | eli. | mean (%) | 0.153 | 45.540 | 25.122 | 5.853 | 0.880 | 0.283 |
| | | SD (%) | 0.000 | 19.976 | 31.655 | 7.124 | 1.642 | 0.174 |
| | | p value | 0.6969 | 0.0007 | 0.8095 | 0.0371 | 0.4475 | 0.1797 |
| Clc3 | veh. | mean (%) | 0.826 | 16.196 | 0.217 | 33.737 | 0.328 | 0.153 |
| | | SD (%) | 1.007 | 5.451 | 0.224 | 22.547 | 0.211 | 0.104 |
| | eli. | mean (%) | 1.181 | 45.552 | 2.409 | 4.849 | 0.349 | 0.580 |
| | | SD (%) | 1.444 | 13.311 | 3.394 | 2.765 | 0.128 | 0.361 |
| | | p value | 0.6259 | 0.0004 | 0.0879 | 0.0033 | 0.7909 | 0.0028 |

**Extended Data Table 6. Quantifications of IHC markers in NB.**

| | | PDX model | MNA NB_2 | MNA NB_3 | MNA NB_4 | MNA NB_5 | NMNA NB_1 | NMNA NB_2 | NMNA NB_3 | NMNA NB_4 |
|---|---|---|---|---|---|---|---|---|---|---|
| Ki-67 | veh. | mean (%) | 92.557 | 87.848 | 67.161 | 31.845 | 6.299 | 64.243 | 45.709 | 63.861 |
| | | SD (%) | 4.386 | 8.549 | 9.169 | 8.726 | 3.544 | 9.930 | 7.811 | 5.695 |
| | eli. | mean (%) | 91.458 | 57.289 | 97.911 | 9.601 | 7.254 | 36.932 | 5.095 | 47.688 |
| | | SD (%) | 7.053 | 5.745 | 0.810 | 4.849 | 3.951 | 9.034 | 1.737 | 10.948 |
| | | p value | 0.6665 | 2E-05 | 3E-06 | 4E-05 | 0.5509 | 0.0001 | 9E-08 | 0.0009 |
| pHH3 | veh. | mean (%) | 4.733 | 1.258 | 0.731 | 0.004 | 0.094 | 2.635 | 1.734 | 3.694 |
| | | SD (%) | 1.225 | 0.890 | 0.350 | 0.013 | 0.095 | 0.905 | 0.790 | 0.630 |
| | eli. | mean (%) | 2.915 | 2.216 | 2.295 | 0.019 | 0.000 | 1.944 | 0.340 | 2.979 |
| | | SD (%) | 1.178 | 0.926 | 0.647 | 0.039 | 0.000 | 0.789 | 0.190 | 1.339 |
| | | p value | 0.0177 | 0.0894 | 0.0003 | 0.3347 | 0.0156 | 0.1392 | 0.0016 | 0.2307 |
| yH2AX | veh. | mean (%) | 19.178 | 20.614 | 2.271 | 0.017 | 0.029 | 4.612 | 5.943 | 3.924 |
| | | SD (%) | 9.551 | 19.203 | 1.983 | 0.039 | 0.052 | 5.741 | 6.480 | 1.850 |
| | eli. | mean (%) | 12.486 | 3.892 | 55.798 | 0.274 | 0.396 | 11.043 | 0.402 | 2.366 |
| | | SD (%) | 9.202 | 1.283 | 20.446 | 0.493 | 0.343 | 6.299 | 0.342 | 2.203 |
| | | p value | 0.0524 | 0.0287 | 2E-05 | 0.1289 | 0.0064 | 0.0719 | 0.031 | 0.1325 |
| Clc3 | veh. | mean (%) | 0.370 | 3.395 | 14.873 | 82.329 | 81.113 | 0.105 | 4.739 | 17.518 |
| | | SD (%) | 0.221 | 1.298 | 10.746 | 16.114 | 10.061 | 0.083 | 2.050 | 9.114 |
| | eli. | mean (%) | 0.825 | 2.524 | 11.238 | 15.229 | 95.581 | 1.148 | 11.204 | 24.413 |
| | | SD (%) | 0.345 | 1.222 | 3.806 | 20.552 | 3.828 | 1.694 | 8.635 | 12.183 |
| | | p value | 0.0082 | 0.1952 | 0.4136 | 0.0001 | 0.0042 | 0.0963 | 0.0398 | 0.2408 |
